# EventHorizon: A Foundation Model for Clinical Flow Cytometry

**DOI:** 10.64898/2026.06.18.733197

**Authors:** Mattia Medina Grespan, Muir J. Morrison, Brendan O’Fallon, Jeffrey R. Jacobsen, Ryan C. Shean, Nicholas C. Spies, David P. Ng

**Affiliations:** ARUP Institute for Research and Innovation, ARUP Labs, 500 Chipeta Way, Salt Lake City, 84108, Utah, USA; Department of Pathology, University of Utah, Salt Lake City, 84108 UT, USA

**Author notes:** These authors contributed equally to this work (Mattia Medina Grespan, Muir J. Morrison).

**Keywords:** Flow Cytometry, Self-Supervised Learning, Diagnostic, Machine Learning

## Abstract

Clinical flow cytometry is central to diagnosing hematologic malignancies, but its reliance on manual expert interpretation and fixed panel designs limits scalability and adaptability. To address this, we developed EventHorizon, a self-supervised foundation model that combines a hierarchical, marker-aware transformer with self-distillation pre-training. EventHorizon integrates heterogeneous, multi-tube panels into a unified, specimen-level embedding. Pre-trained without labels on 100 937 routine clinical specimens—comprising over 50 billion cells—the model yields frozen embeddings that enable lightweight classifiers to accurately identify recurrent genetic abnormalities in acute myeloid leukemia (AML), including t(15;17) (AUROC 0.991) and an unexpected signal for *DDX41* mutations (AUROC 0.970). EventHorizon maintained strong performance on a temporally separated 2026 cohort (macro AUROC 0.970 across 28 diagnoses) and demonstrated zero-shot transferability across a five-site B-cell lymphoma cohort (CLL vs. normal AUROC 0.988), the FlowCAP-II AML challenge (AUROC 0.973), and B-ALL measurable residual disease detection (AUROC 0.920 at ≥1% disease burden). Although class ranking transferred reliably across sites, decision thresholds shifted; however, these were rapidly recalibrated using as few as four labeled AML cases. Sensitivity decreased at disease burdens below 0.1%, yet embeddings remained robust to simulated assay perturbations. Overall, EventHorizon offers a reusable, panel-agnostic representation for diagnostic flow cytometry.

## 1. Introduction

Clinical flow cytometry is essential for diagnosing hematologic malignancies, but manual interpretation creates throughput bottlenecks, limits scalability, and introduces inter-operator variability [1, 2]. These challenges are exacerbated by workforce constraints and reimbursement pressures [2–4]. While machine learning methods have been proposed [5–14], they typically rely on either high-level supervision from raw embeddings [5, 8, 9, 11, 12, 14] or low-level supervision from predefined gating [13, 15]. Both paradigms require large labeled datasets, are sensitive to panel design, and struggle to generalize across laboratories due to technical variability [9, 16]. Self-supervised learning (SSL) offers a promising alternative for learning representations from unlabeled data [17, 18]. Existing flow cytometry SSL approaches mainly use specimen-level encoders (often paired with multiple-instance learning) [14, 18, 19] or flexible single-cell encoders [17, 20]. Specimen-level encoders yield clinically useful representations but are bound to fixed panels. Single-cell encoders accommodate variable panels but require post hoc pooling to generate specimen-level representations, limiting clinical utility. Methods like GPCT [21] and FATE [22] attempt to bridge this gap via self-attention across cells, but still suffer from reliance on post hoc pooling [21], limited dataset scale, or task-specific constraints [22].

In this work, we introduce *EventHorizon*, a foundation model for flow cytometry that integrates panel flexibility with direct learning of cell- and specimen-level representations. Building on DINO-style self-distillation [23–26] and recent transformer architectures [18, 20, 22, 27], EventHorizon maps heterogeneous cell collections across variable panels into unified specimen embeddings. This design eliminates technical artifacts and workflow variations while preserving diagnostic biological signal.

## 2 Methods & Data

### 2.1 Datasets

EventHorizon was pre-trained on 100 937 clinical specimens collected during routine practice at ARUP Laboratories from mid-2019 through 2025, reflecting the natural distribution of cases in a reference laboratory (Supplemental Table S2). Each specimen was evaluated with panel combinations driven by specimen type, patient history, and suspected diagnosis. Peripheral blood and bone marrow aspirates always receive B-cell, T-cell, and myeloid screening tubes (hereafter, triage panel), with marrows additionally receiving a plasma cell tube, and add-on tubes are ordered based on initial findings. In total, the dataset contains 59 unique markers across 95 antibody–fluorophore pairs (Supplemental Tables S3), all acquired on six Beckman Coulter Navios cytometers. Pre-processing is described in Supplemental Section 1.2.

To evaluate embedding quality, diagnostic labels (50 multi-label categories across 10 disease groups, with specimen metadata and uncertainty annotations, see Supplemental Table S2) were extracted from final flow cytometry reports using a large language model (Qwen3.5-122B-A10B-FP8 [28]) in a two-stage pipeline with deterministic post-processing. The schema extends a previously validated BioMedBERT-based system [10], with high concordance on overlapping categories (Supplemental Section 1.3).

### 2.2 Model Architecture

EventHorizon uses a 2-tier hierarchical Transformer encoder designed to combine the marker flexibility of cell-level encoders [17, 20, 21] with the specimen-level representations produced by fixed-panel models [14, 18, 19]. As shown in Figure 1, marker identity and expression are first jointly encoded for each cell, allowing cells measured with different marker panels to be represented in a common latent space. A first “marker self-attention” block integrates information among markers within each cell to produce cell-level representations, after which a second “cell self-attention” block integrates cells across all included panels to generate a single specimen-level representation. This design allows EventHorizon to jointly encode multiple heterogeneous panels from a specimen without requiring a fixed marker configuration. See SI Section 1.5 for architectural and implementation details.

**Fig 1:**
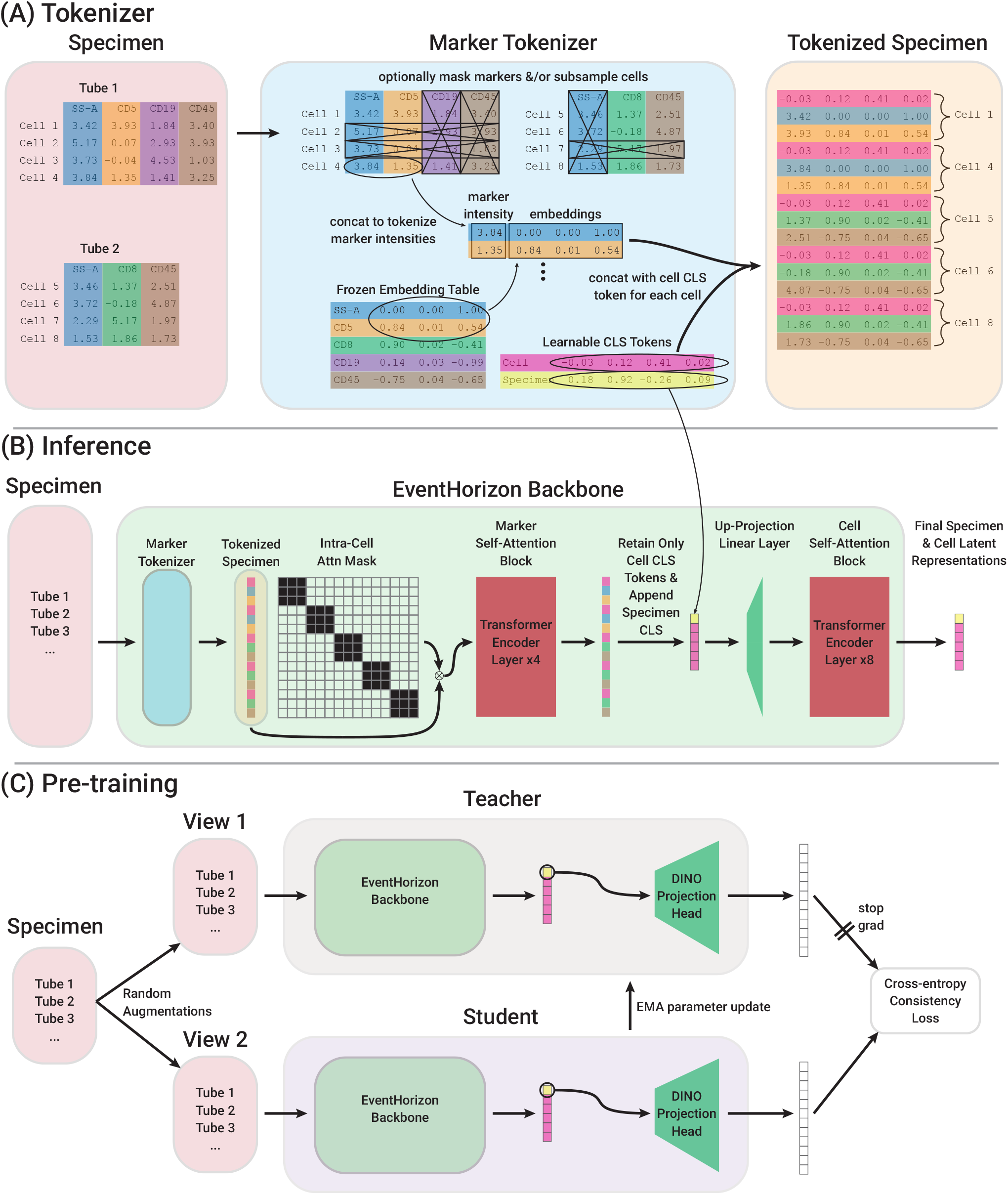
Overview of EventHorizon. (A) The tokenizer design which enables our hierarchical encoder architecture. This tokenization strategy allows us to seamlessly flatten data from multiple marker panels from a single specimen into a single “bag” of tokens. (B) The 2-stage backbone of our model in inference mode. In the first stage, masking allows only intra-cell self-attention, while in the second stage, all cells from a specimen attend to each other. This enables marker/panel flexibility in the first stage combined with inter-cell self-attention and specimen-level representations in the second. (C) Self-supervised pre-training workflow, similar to DINO and other joint-embedding architectures, but with flow cytometry-specific augmentations (see Sections 2.2, 2.3 and Supplemental Information, Section 1.6 for details).

### 2.3 Training Strategy

EventHorizon follows the DinoFlow framework [18], which builds on DINO [23] and DINOv2 [24]. A projection head is appended to the backbone during pre-training [29], and identical student and teacher networks process multiple augmented views of each specimen (Figure 1). A cross-entropy objective aligns representations across views, while an auxiliary KDE regularizer encourages uniform feature spread. Teacher output centering is used to mitigate mode collapse. The student is optimized by gradient descent and the teacher is updated as an exponential moving average of the student. After pre-training, the projection head is discarded and the teacher backbone is retained for downstream applications.

The main novelty in our training strategy lies in the design of our augmentations, since it is far from obvious how to translate the image augmentations from image SSL frameworks such as DINO [23] and DINOv2 [24] to flow cytometry data. Broadly, we combine flow cytometry-specific augmentations introduced by DinoFlow [18], masking inspired by prior flow cytometry SSL strategies [17, 19–22], perturbations to the spillover matrix, and a novel flow cytometry-specific cropping strategy. See SI Section 1.6 for details and hyperparameters.

### 2.4 Embedding generation for downstream evaluation

For downstream analyses, embeddings were generated using the frozen EventHorizon teacher backbone. For each specimen, events from the tubes used in a given evaluation were jointly passed through EventHorizon, and the resulting specimen-level representation was used as the specimen embedding. Unless otherwise specified, downstream embeddings were generated using a total event budget of 217 (131 072) events per specimen, split across the tubes included in the corresponding analysis. Evaluations used either all tubes retained for a specimen after preprocessing (hereafter, “all tubes”), a predefined subset of tubes, or a single tube, as specified for each analysis. For external datasets, marker names were harmonized to the EventHorizon vocabulary where possible, with unrecognized channels handled as described in SI, Section 1.5.

### 2.5 Latent-space graph analysis

To assess the organization of the EventHorizon latent space, we analyzed peripheral blood and bone marrow specimens containing the triage panel tubes described in Section 2.1. Cosine *k*-nearest-neighbor graphs were constructed from specimen embeddings generated using all tubes, and graph structure was evaluated using modularity and Leiden community detection [30, 31]. We evaluated organization with respect to diagnosis and tube-panel composition using stratified permutation tests and community purity. The primary analysis used *k* = 5 and Leiden resolution 1.0.

We additionally compared tube-level EventHorizon and DinoFlow embeddings from the triage panel tubes with respect to diagnosis, tube type, and cytometer ID. See SI Section 1.8 for graph construction, permutation, community-purity, and parameter-sensitivity procedures.

### 2.6 Comparison models

For the triage evaluation, we compared EventHorizon with DinoFlow [18] and an adapted Farthest Point Sampling SetTransformer (ST-FPS) [32] using the triage panel tubes. EventHorizon and DinoFlow were evaluated using frozen self-supervised representations, whereas ST-FPS was trained end-to-end from raw ell measurements using supervised diagnostic labels. See SI Section 1.7 for model adaptation, representation construction, training, and hyperparameter selection.

## 3 Results

### 3.1 AML Molecular Subclassification

To evaluate the biological significance of the embedding space, we obtained molecular and cytogenetic data within a two-week window for all acute leukemias (≥15% blasts, excluding MRD studies). This yielded 2 589 cases: 2 337 acute myeloid leukemias, 55 MPAL/AUL, 163 B-ALL, and 34 T-ALL. EventHorizon embeddings were generated from all tubes per specimen (Section 2.4). For each abnormality, we evaluated logistic regression performance using 5-fold cross-validation, reporting AUROC and AUPRC (to account for class imbalance) with 95% confidence intervals from 1 000 bootstrap resamples.

As expected, these embeddings recapitulated the known phenotypic landscape of acute leukemia (Figure 2A) and its established phenotype–genotype correlates (Figure 2B). Reflecting their distinctive immunophenotypes, high classification performance was observed across key entities:

- **t(15;17) / APL** (*n* = 5): AUROC 0.991 [0.986–0.996], AUPRC 0.810 [0.692–0.902]
- **t(8;21) *RUNX1::RUNX1T1*** (*n* = 13): AUROC 0.981 [0.961–0.996], AUPRC 0.613 [0.350–0.832]
- **inv(16)/t(16;16)** (*n* = 22): AUROC 0.967 [0.914–0.996], AUPRC 0.674 [0.480–0.875]
- **Cytogenetically normal *NPM1* -mutated** (*n* = 166): AUROC 0.887 [0.857–0.917], AUPRC 0.525 [0.444–0.609]
- **MDS-related gene mutations** (*n* = 979): AUROC 0.764 [0.743–0.785], AUPRC 0.728 [0.698–0.758]

Notably, 50 cases with *DDX41* mutations achieved an AUROC of 0.970 [0.954–0.982] (AUPRC 0.537 [0.399–0.668]). This performance exceeds that of the well-described *NPM1* immunophenotype, suggesting a previously unrecognized immunophenotypic signature in *DDX41* -mutated AML.

A *K*-nearest neighbors classifier (*K* = 25, cosine distance) performed significantly worse on rare abnormalities (e.g., *RUNX1::RUNX1T1* AUROC 0.789; *DDX41* AUROC 0.892), as expected when positive class size approaches or falls below *K*. Performance metrics for additional molecular and cytogenetic targets are provided in SI Table S5.

**Fig 2:**
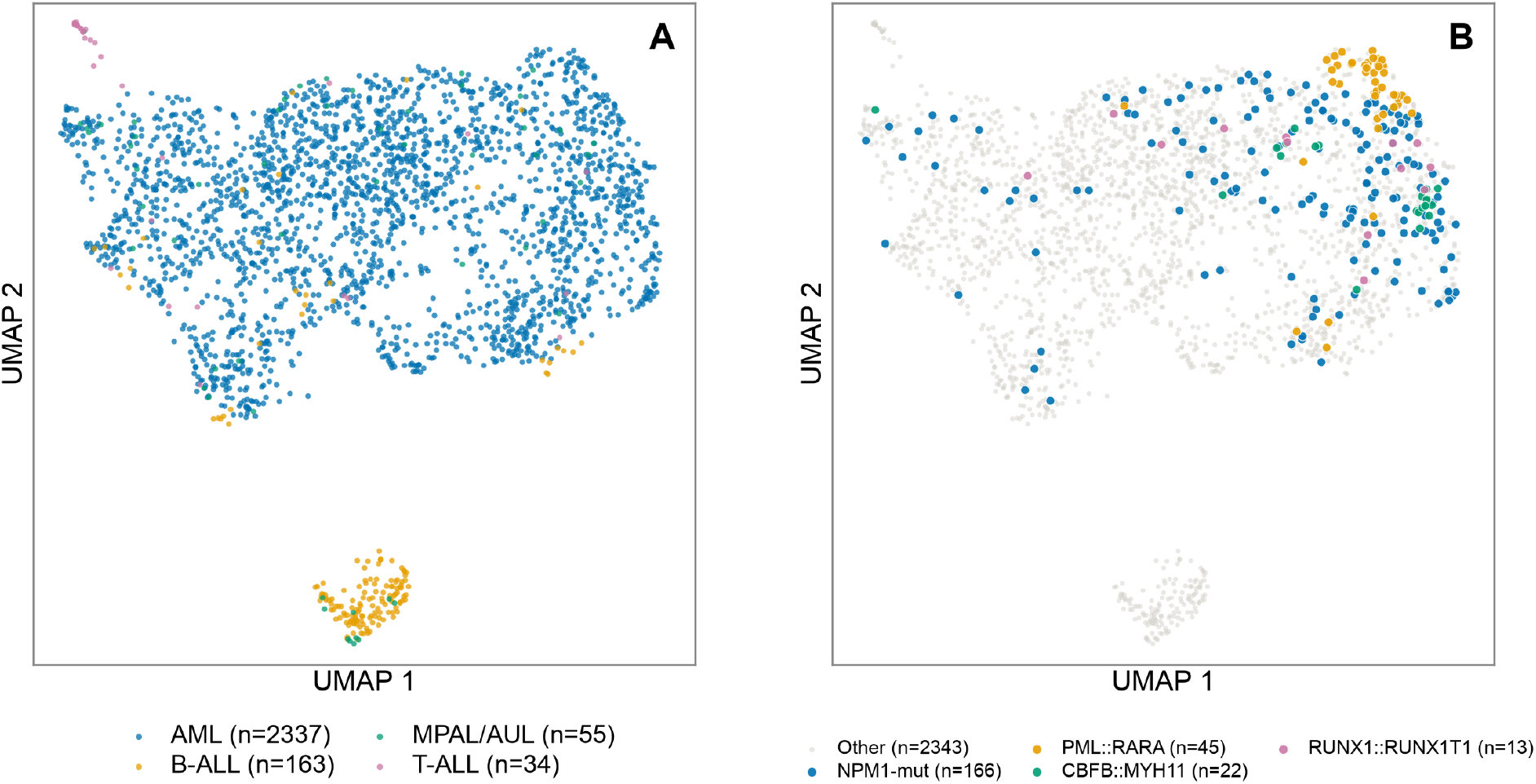
UMAP projection of full case embeddings for the acute leukemia cohort, colored by (A) diagnostic subtype and (B) WHO-defined recurrent genetic abnormality. In (B), cases are colored by a subset of mutually exclusive molecular subtypes, include NPM1 mutated without cytogenetic abnormalities

A *K*-nearest neighbors classifier (*K* = 25, cosine distance) performed significantly worse on rare abnormalities (e.g., *RUNX1::RUNX1T1* AUROC 0.789; *DDX41* AUROC 0.892), as expected when positive class size approaches or falls below *K*. Performance metrics for additional molecular and cytogenetic targets are provided in SI Table S5.

### 3.2 Temporal data split performance

To evaluate clinical classification performance, we embedded 110 041 cases (2019–2025; Section 2.4), excluding proficiency, development, and validation samples. We used an 85/15 temporal train/test split (93 535 train/16 506 test) to train five classifiers on clinically derived labels: a multilayer perceptron (MLP), logistic regression, XGBoost, random forest, and *k*-nearest neighbors (*k*-NN).

Models were evaluated without tuning on an independent prospective cohort of 10 901 sequential cases collected after January 1, 2026. Classification metrics were macro-averaged across 28 diagnostic labels with *≥* 10 positive cases, using decision thresholds fixed at each label’s *F*1-maximizing point on the test set: **Multilayer Perceptron:** AUROC 0.970 [0.965–0.974], AUPRC 0.486 [0.465–0.507], *F*1 0.468 [0.447–0.487] **Logistic Regression:** AUROC 0.966 [0.960–0.971], AUPRC 0.440 [0.423–0.458], *F*1 0.442 [0.425–0.459] **XGBoost:** AUROC 0.961 [0.956–0.965], AUPRC 0.452 [0.431–0.474], *F*1 0.443 [0.423–0.463] **Random Forest:** AUROC 0.944 [0.935–0.953], AUPRC 0.427 [0.408–0.449], *F*1 0.429 [0.411–0.449] *k***-Nearest Neighbors:** AUROC 0.886 [0.874–0.898], AUPRC 0.401 [0.380–0.421], *F*1 0.408 [0.387–0.429] Full per-label AUPRC and *F*1 performance, with 95% confidence intervals, is provided in SI Table S6.

### 3.3 Zero-shot Evaluation of B Cell Non-Hodgkin Lymphomas

To evaluate generalization to out-of-distribution datasets, we mapped our diagnostic labels to the schema of a multi-center German B-NHL dataset (*n* = 36 542 across 5 sites; Table 1) [9]. Substantial domain shifts existed between cohorts, including missing antibody markers (IgM, sCD79b, CD43, FMC7), restriction to mature B-cell neoplasms, and specimen selection limited to bone marrow and peripheral blood (unlike our multi-tissue pretraining set).

**Table 1:** ARUP-to-German label mapping based on panel design and clinical-practice characteristics as determined by an expert hematopathologist. Labels were scored with an ARUP-trained *K*-NN classifier (*K* = 25, cosine distance) on German embeddings at 217 events. *F*1 uses each mapping’s own *F*1-maximizing threshold on the German cohort. Prevalence is out of 36 542 German specimens.

| ARUP source | German target(s) | $n_{\text{pos}}$ | Prev. (%) | AUROC | AUPRC | $F_1$ |
| --- | --- | --- | --- | --- | --- | --- |
| CLL | MBL+MLUS+<br>CLL+atypbcll+PL | 10 400 | 28.46 | 0.970 [0.968–0.972] | 0.951 [0.948–0.955] | 0.921 [0.916–0.924] |
| 5+BNHL | MCL | 549 | 1.50 | 0.795 [0.775–0.815] | 0.071 [0.060–0.085] | 0.151 [0.129–0.174] |
| HCL_variant | HCLv | 108 | 0.30 | 0.788 [0.742–0.836] | 0.038 [0.018–0.071] | 0.094 [0.044–0.148] |
| 10+BNHL | FL+DLBCL | 503 | 1.38 | 0.775 [0.751–0.798] | 0.085 [0.070–0.108] | 0.161 [0.133–0.192] |
| PC_differentiation | LPL | 890 | 2.44 | 0.761 [0.743–0.778] | 0.129 [0.112–0.151] | 0.229 [0.210–0.251] |
| HCL | HCL | 587 | 1.61 | 0.754 [0.731–0.774] | 0.251 [0.212–0.289] | 0.317 [0.278–0.353] |
| 5-10-BNHL | MZL | 1 240 | 3.39 | 0.556 [0.544–0.570] | 0.039 [0.037–0.043] | 0.086 [0.079–0.094] |

Despite these shifts, zero-shot transfer performance was strong for well-represented classes. For the binary task of distinguishing chronic lymphocytic leukemia (CLL) from normal specimens (*n* = 29 623), an ARUP- trained kNN classifier (*K* = 25, cosine distance) achieved an AUROC of 0.988 [0.986–0.990] (accuracy 0.969, sensitivity 0.973, specificity 0.968) using an ARUP-fixed threshold, closely matching internal performance (AUROC 0.997 [0.995–0.999]). Performance on the broader CLL-adjacent cluster is detailed in Table 1.

Given label noise and schema discrepancies, we analyzed nearest-neighbor geometry in the embedding space to assess biological alignment. The single nearest ARUP match for a German MBL or MLUS specimen carried the ARUP CLL label in 43–66% of cases (5.1×–7.8× enrichment over its 8.4% ARUP baseline). For hairy cell leukemia (HCL), 28.8% of German cases had an ARUP HCL match as their nearest neighbor—a 129.3-fold enrichment over baseline prevalence (0.23%) and the highest enrichment observed across all labels. Conversely, HCL variant (HCLv) demonstrated a distinct pattern: ARUP’s HCLv was only the sixth most frequent nearest match (7.4%), trailing the broader 10+BNHL category (29.6%), suggesting potential differences in diagnostic classification criteria between cohorts.

### 3.4 Zero-shot Evaluation on the FlowCAP-II AML Dataset

As a further evaluation of EventHorizon on an out-of-distribution dataset, we evaluated the 359 cases (43 AML, 316 normal) of the seminal FlowCAP-II AML challenge [15] zero-shot, that is, with classifiers trained only on ARUP embeddings and no FlowCAP data. Because the FlowCAP annotations are limited to AML versus normal, and we cannot be certain whether FlowCAP normal cases are strictly normal or simply not AML, we evaluated the AML-vs-normal task, whose label definition most closely mirrors the FlowCAP annotation. Classifiers were trained on 85% of the ARUP AML-vs-normal cases (5 800 AML/35 830 normal in total), and the remaining 15% was held out to select the classification threshold (F1-maximizing). The kNN classifier (K=25, cosine distance) achieved an AUROC of 0.973 [95% CI 0.942,0.993] and an AUPRC of 0.812 [0.695,0.922] on the 359 FlowCAP cases. On the original FlowCAP-II test partition (20 AML, 160 normal), the kNN achieved an AUROC of 0.993 [0.984,1.000]. Interestingly, case 116 was misclassified as normal by the kNN classifier (AML probability 0.12), and on manual review by two expert hematopathologists (DPN and JRJ), this was deemed a mis-annotation, a finding previously reported by Biehl et al. [33]. Ranking performance transferred well, but the classification threshold did not. Using the ARUP-derived threshold, the kNN achieved a sensitivity of 0.977 [0.923,1.000] but a specificity of only 0.854 [0.814,0.890] (F1 0.641 [0.547,0.732]). Recalibrating the threshold required very few labeled FlowCAP cases from the train partition: 4 AML and 27 normal cases restored the kNN F1 score on the test partition from 0.66 [0.53,0.77] to 0.90 [0.79,0.97] and specificity from 0.87 to 0.99, with further gains up to an F1 of 0.95 [0.88,1.00] when all 23 AML cases were used (Figure 3).

**Fig 3:**
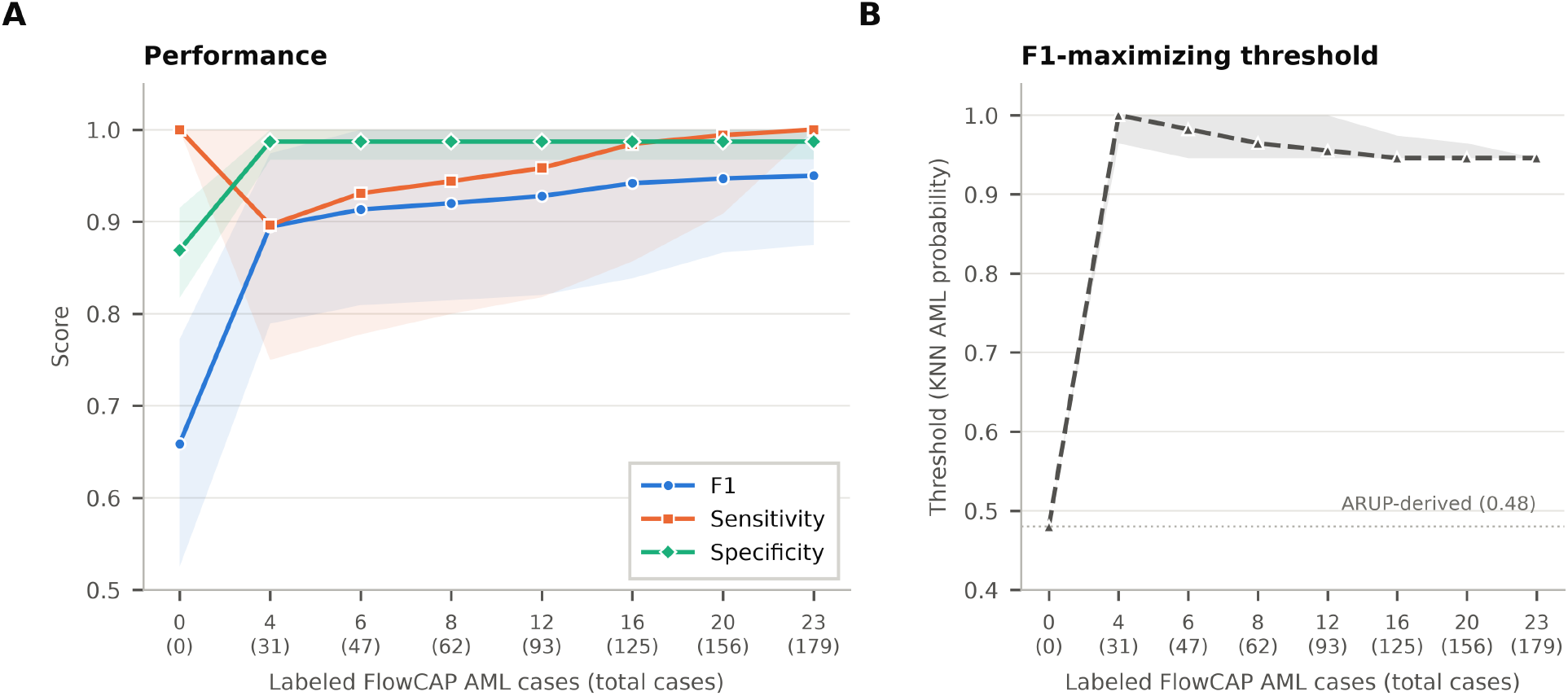
Few-label threshold recalibration on FlowCAP-II. (**A**) F1, sensitivity, and specificity of the ARUP-trained kNN on the FlowCAP-II test partition as a function of the number of labeled FlowCAP training-partition AML cases (total cases in parentheses, normals in pool proportion) used to re-select the F1-maximizing threshold; 0 denotes the ARUP-derived threshold. (**B**) The selected threshold (dotted line: ARUP-derived, 0.48). Lines are means (A) or medians (B) over 30 random adaptation draws; bands are 2.5– 97.5 percentiles (A: nested bootstrap over 100 test-set resamples per draw; B: across draws).

### 3.5 Zero-shot Evaluation on B-ALL: Classification and Regression

To further characterize generalizability, we tested the model on held-out Children’s Oncology Group (COG) B-ALL measurable residual disease (MRD) samples. Day 8 single-tube panels were downsampled to 217 events. Day 29 three-tube panels excluded the final SYTO 16 tube, downsampling the remaining two tubes to 216 events each while omitting markers absent from the pre-training vocabulary (CD9, CD13/CD33 stack). Applying our multi-label MLP classifier to the COG embeddings showed performance scaling directly with disease burden: AUROC increased from 0.694 across all 420 MRD-positive cases (*n* = 921 negatives) to 0.696 at ≥0.01% (*n* = 414), 0.773 at ≥0.1% (*n* = 287), and 0.920 at ≥1.0% (*n* = 132), with AUPRC following the same trend (0.569 →0.567 →0.597 →0.721).

To evaluate disease burden estimation, we trained a ridge regressor on log-transformed report burdens from 776 pre-training adult de novo B-ALL cases (median 50%, range 0.01%–99%). This training set was out-of-distribution for the COG testing cohort (median burden 0.25%), which differs in clinical context (pediatric post-treatment MRD vs. adult diagnostic/de novo), and target disease range. Consequently, regressor performance hinged on training distribution overlap (Figure 4). Above 1% disease burden (*n* = 132), where the COG cases overlapped with the adult training set, the model performed strongly and remained well-calibrated (*R*2 = 0.605, *ρ* = 0.760, median bias 1.17*×*). In contrast, for out-of-distribution cases below 1% (*n* = 288), performance collapsed entirely (*R*2 = *™*4.14, *ρ* = 0.235, median bias 11.6*×*).

**Fig 4:**
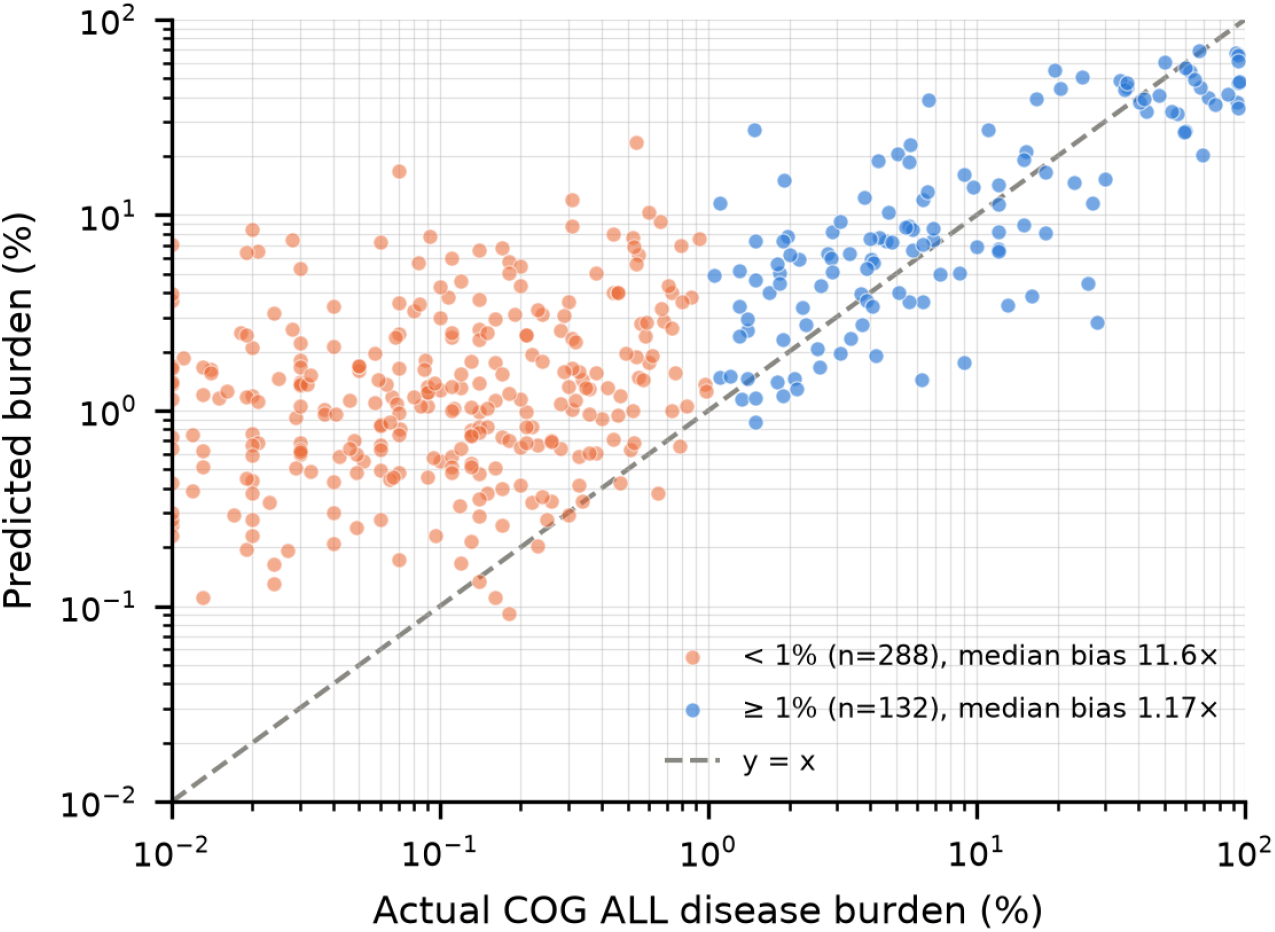
Results of ridge regressor trained on B-ALL disease burdens and EventHorizon Embeddings. Predictions vs. reported disease burden in the COG ALL external validation cohort (log–log scale).

### 3.6 Zero-shot Analysis of Spectral Flow Data

We evaluated EventHorizon on 28 residual clinical specimens run on a developmental 37-color spectral panel against diagnoses from a blinded hematopathologist (Supplemental Section 2.2). Using the top label among 25 nearest ARUP neighbors (excluding matched Navios runs), top-1 accuracy reached 46% (13/28) and top-neighbor coverage reached 71% (20/28; Supplemental Figure S1). Agreement was highest for myeloid neoplasms (6/7) and lowest for B-cell neoplasms (0/5).

### 3.7 Sensitivity Analysis on Embedding Events

To assess how embedding event count affects disease-detection sensitivity across the CLL/SLL–CD5+MBL disease continuum, we evaluated 214-, 216-, and 217-event embeddings on a balanced binary cohort (9 701 CLL or CD5+MBL cases; 9 701 normal controls sampled to match positive specimen types). Performance was evaluated using stratified five-fold cross-validation with 5-nearest-neighbor cosine similarity.

To isolate the effect of event count in the reference set versus the query specimen, we compared extreme combinations of 214- and 217-event embeddings (Figure 5). Among 9 522 positive cases with reportable disease burden, performance differences were confined to low-burden specimens (*>* 0–0.1%). In this regime, increasing the query event count from 214 to 217 improved sensitivity, whereas increasing the reference event count alone provided minimal benefit. Above 0.1% burden, sensitivity was comparable across all conditions and rapidly approached a ceiling. These findings suggest that at very low disease burdens, performance is improved primarily from acquiring additional events for the query specimen at inference, rather than for the kNN reference/training embeddings.

**Fig 5:**
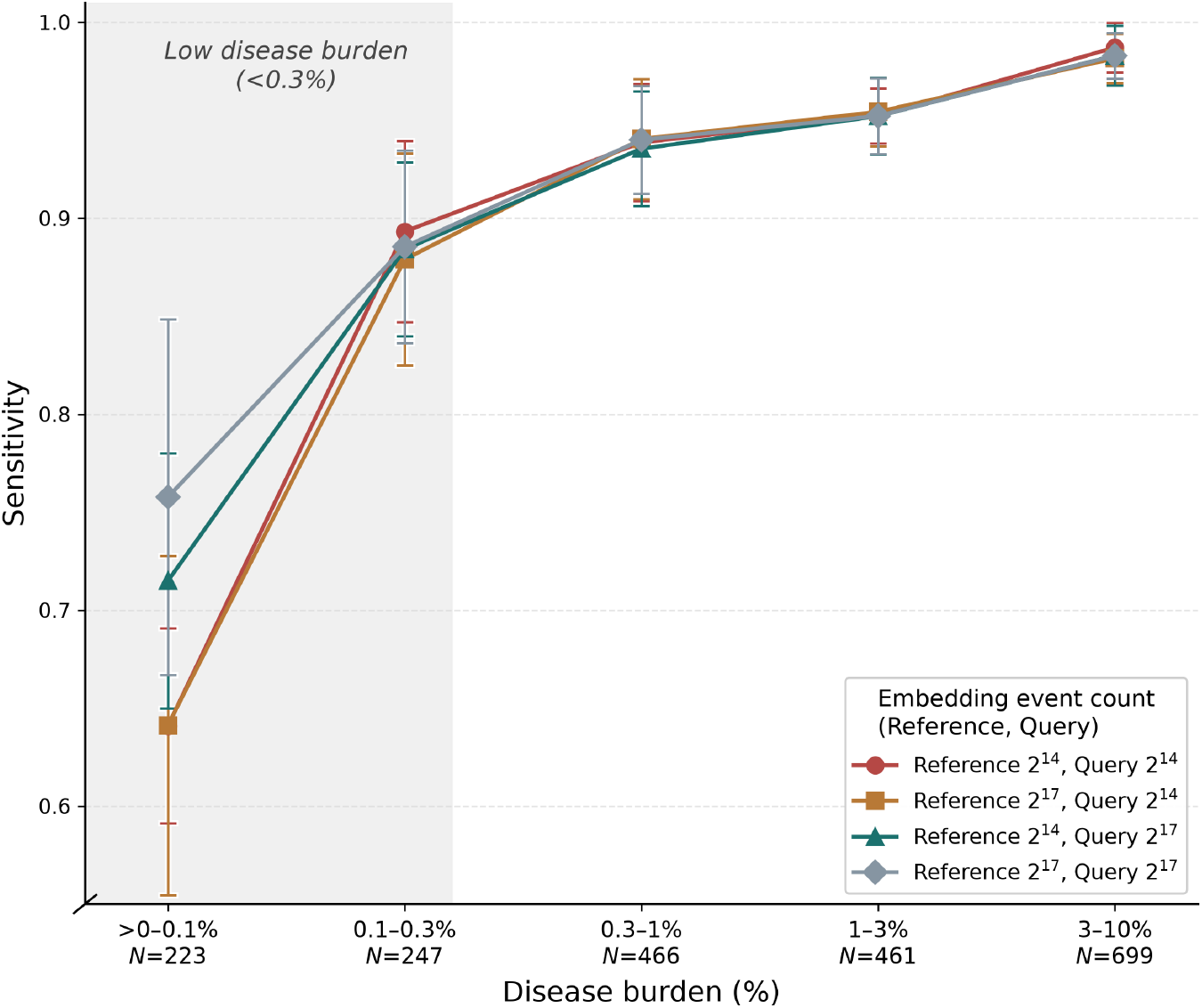
Effect of embedding event count on sensitivity across the CLL/SLL–CD5+MBL disease-burden continuum. Sensitivity was evaluated using 5-nearest-neighbor classification with cosine similarity and five-fold cross-validation. Lines show mean sensitivity across folds and error bars indicate ±1 standard deviation. The four conditions compare 214 and 217 event embeddings for the kNN reference set and held-out query specimens. The shaded region indicates low disease burden (*<* 0.3%). The number of positive specimens in each disease-burden bin is shown below the x-axis.

### 3.8 Perturbation Analysis

To evaluate embedding robustness to laboratory perturbations, a held-out 2026 cohort (*n* = 6 579) was reembedded under increasing scale, shift, and spillover perturbations across three independent trials using established protocols [18]. Cases were embedded at 214 events due to computational constraints. Classification performance was evaluated using the top-performing MLP classifier without retuning, retaining decision thresholds fixed on the 2019–2025 test set.

Consistent with prior findings [18], embeddings demonstrated marked resilience to simulated noise, preserving performance at perturbations up to 3*σ*(Figure 6). Notably, scaling errors induced a larger performance dropoff than shift or spillover perturbations, contrasting with trends reported in DinoFlow.

**Fig 6:**
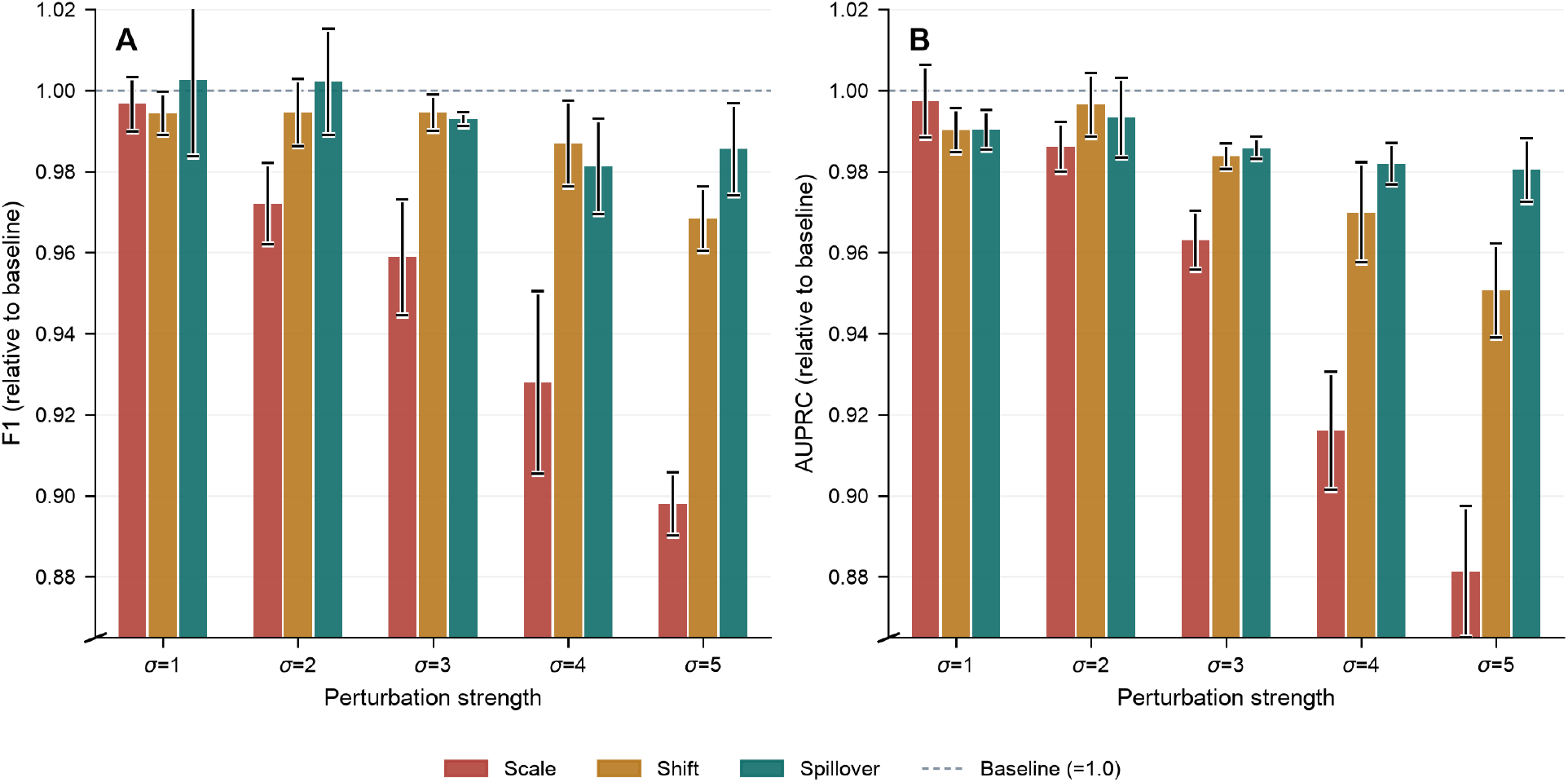
Embedding robustness across simulated data perturbations. (**A**) Macro *F*1 score (unweighted mean across 36 labels with ≥10 positive cases). (**B**) Macro area under the precision–recall curve (AUPRC). Values are relative to baseline (*σ*= 0; dashed line at 1.0); error bars represent ±1*σ*across three random seeds. The *y*-axis is truncated to highlight differences between conditions.

### 3.9 Comparison with other models

We evaluated diagnostic classification on specimens containing the triage panel. Models were developed using 78 747 historical specimens from 2019–2025 and evaluated on a strict 2026 holdout cohort of 8 034 specimens not used for pre-training or supervised model development, across 33 diagnostic labels with *≥* 10 holdout occurrences (SI Table S2). ST-FPS achieved the highest aggregate *F*1 and AUPRC, followed by EventHorizon using all tubes, EventHorizon using triage tubes only, and DinoFlow (Figure 7). Per-label performance was more variable: frozen EventHorizon embeddings performed comparably to ST-FPS for common diagnoses including NORMAL, CLL, and AML, and were competitive with or exceeded ST-FPS for several individual diagnoses. ST-FPS showed larger advantages only for phenotypes requiring fine-grained event-level resolution, such as CD4/CD8 ratio abnormalities (full per-label results in SI Figure S2). Results using *k*-nearest-neighbor probes are provided in SI Figure S3.

**Fig 7:**
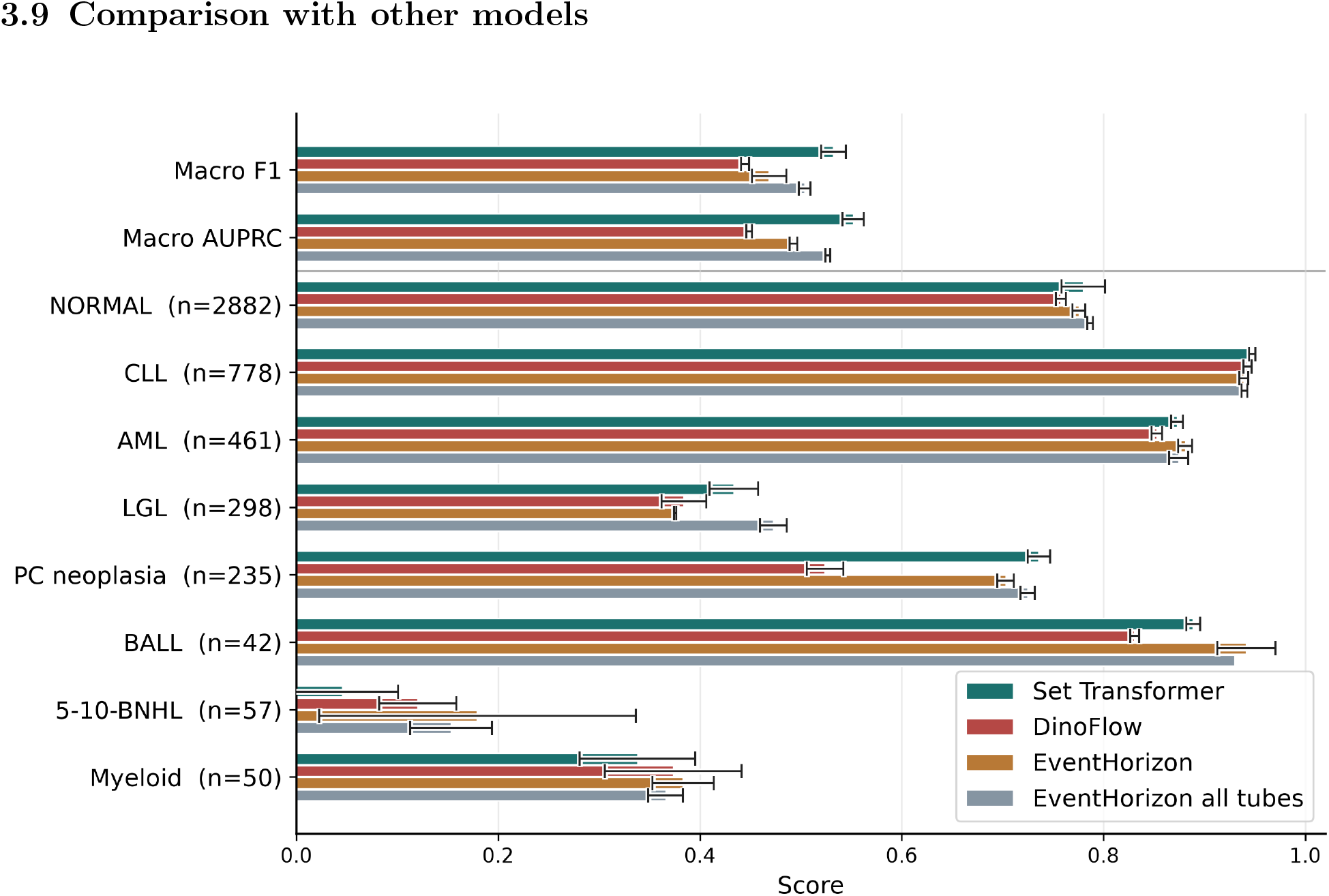
Binary diagnostic performance on the triage screening evaluation. (Top) Macro *F*1 and macro AUPRC across the full set of 33 evaluated diagnoses. (Bottom) Per-label *F*1 for selected diagnoses (mean ± SD across three random seeds). EventHorizon and DinoFlow rely on frozen self-supervised specimen embeddings evaluated with MLP probes; ST-FPS serves as an end-to-end supervised baseline.

### 3.10 Biological and technical organization of the latent space

To determine whether EventHorizon specimen embeddings encode diagnostic structure beyond tube-panel composition (which correlates with diagnosis due to clinical workup workflows), we performed graph-based latent space analysis (Section 2.5).

At *k* = 5, diagnosis modularity reached 0.39 versus a stratified permutation mean of 0.06 (*z* = 333.2), whereas tube-panel signature modularity was 0.22 (permutation mean 0.13, *z* = 98.6). Leiden community analysis showed consistent trends: at resolution 1.0, diagnosis purity was 0.89 (majority-label baseline 0.67), while tube-panel signature purity was 0.78 (baseline 0.71). These findings were robust across varying values of *k* and Leiden resolutions (SI Figures S5–S8).

Comparing tube-level latent spaces, EventHorizon demonstrated stronger diagnostic clustering than DinoFlow (modularity 0.39 vs. 0.30; purity 0.86 vs. 0.83). Conversely, DinoFlow organized predominantly by tube type, exhibiting markedly higher tube-type modularity (0.67 vs. 0.41) and purity (1.00 vs. 0.34). Cytometer ID structure was minimal and comparable across both models (modularity 0.14; purity 0.25 for EventHorizon vs. 0.29 for DinoFlow; SI Figures S9–S10).

Overall, EventHorizon retains strong diagnostic structure while suppressing tube-identity effects compared to DinoFlow; cytometer-associated batch effects remain minimal in both models.

## 4 Discussion

A central question for any self-supervised model is whether its encoding reflects disease biology rather than technical artifacts of the assay. Without any genetic information during pretraining, EventHorizon recovered genetic abnormalities with distinctive immunophenotypes using only a linear classifier, and these results compare favorably with task-specific attention-based models trained on a two-tube panel [14], although differences in cohorts and case selection preclude direct comparison. Conversely, mutations without a characteristic immunophenotype, such as *DNMT3A* and *TET2*, were only modestly predictable. This gradient follows established phenotype–genotype relationships rather than the uniform performance expected from a technical artifact. The latent-space graph analysis (Section 3.10) supports the same conclusion, since diagnosis was far more organized in the embedding than tube-panel composition, which correlates with diagnosis because add-on panels are selected during clinical workup.

The strong signal for *DDX41* -mutated AML, exceeding that of *NPM1*, was unexpected, as no immunophenotype has been reported for this entity. *DDX41* -mutated myeloid neoplasms are clinically distinctive, frequently arising in older men with hypocellular marrows and low blast counts [34], so the embedding may be capturing these specimen-level features rather than a mutation-specific phenotype. Given only 50 cases, we regard this as hypothesis-generating and plan to validate it in an external cohort and to distinguish germline from somatic variants.

The representation generalized across time and sites without retraining: classifiers trained on 2019–2025 data maintained performance on a 2026 cohort, and ARUP-trained models transferred zero-shot to three external datasets with distinct panels and case mixes. While performance dipped where diagnostic criteria differed across institutions, concordant nearest-neighbor enrichment indicates that underlying biological signals were preserved. Class ranking transferred more reliably than calibration; although the ARUP threshold failed on FlowCAP-II, fine-tuning with just 4 AML and 27 normal cases restored performance (Figure 3). Adopting sites can thus recalibrate operating thresholds locally rather than retraining the model. Finally, zero-shot evaluation on an unseen spectral panel showed partial transfer–myeloid specimens retrieved relevant neighbors, whereas entities requiring pretraining-absent markers performed poorly (Supplemental Figure S1). We are developing marker-aware tokenization which may better facilitate spectral integration via light adaptation or additional pretraining.

Performance depended strongly on disease burden. Sensitivity fell substantially below 0.1% burden in the CLL/MBL continuum and in COG B-ALL, and increasing the events used to embed the query specimen, but not the reference set, improved it (Figure 5). The limiting factor at low burden is therefore how completely a rare population is sampled at inference, not the quality of the representation. At 217 events per specimen, a population at 0.01% is represented by only about 13 events, whereas MRD protocols typically acquire *>* 106 events per tube. EventHorizon should thus be regarded as a diagnostic and screening representation rather than an MRD assay.

Robustness to routine assay variation is a key operational requirement, since instrument drift, detector voltage changes, compensation errors, and reagent lot changes alter measured intensities. Classification performance was largely preserved under simulated scale, shift, and spillover perturbations up to three times the within-laboratory standard deviation, without retraining classifiers or adjusting thresholds (Figure 6). This is partly by design, as the perturbations resemble the pretraining augmentations. Scaling errors were the most damaging, in contrast to DinoFlow [18]. This robustness reflects technical variation within one laboratory over time and should not be confused with invariance across sites, where differences in panels, preparation, and populations were larger.

Frozen EventHorizon embeddings with a lightweight probe approached a fully supervised end-to-end model (ST-FPS [22]) on common diagnoses, though end-to-end training remained stronger for phenotypes defined by fine cell-set geometry, such as abnormal CD4:CD8 ratios. The advantage of a general-purpose representation is reuse at a fraction of the cost. At 6.5 million parameters, EventHorizon is small for a foundation model, a size set by compute rather than design. Memory limits were most severe in pretraining, where the batch, augmented views, student and teacher networks, and saved activations must coexist, restricting training views to 1 000–8 000 cells per tube. That the model nonetheless performs well on far larger inputs, and on single tubes, suggests that the representation extrapolates beyond its training regime. Self-attention across cells also remains quadratic in compute at inference. It is therefore unclear whether performance is bounded by data, model capacity, or context length, and 100 000 specimens, although comprising over 50 billion cells, is modest for self-supervised pretraining. Gains in low-burden sensitivity with longer inference contexts suggest that input size is at least one bottleneck, and these results are likely a lower bound for the approach.

We view EventHorizon as an assistive tool rather than a replacement for expert interpretation. The most promising clinical use is case-based decision support: for a difficult specimen, the model can retrieve nearest neighbors in embedding space with their historical sign-out diagnoses, giving pathologists a record of how similar cases were handled across time and institutions. A second use is near-real-time quality control, running the model in the window around sign-out to flag discrepancies with the draft diagnosis, although this will require substantial engineering and workflow integration. Gains in triage, where real-time systems already perform well [10], are likely to be modest. For research, the same representation can identify immunophenotypically similar cohorts across archives, as the *DDX41* signal suggests, and pool data across laboratories with different panels. Any clinical deployment will require local validation, particularly on ambiguous cases. This study has important limitations. Pretraining and most evaluations used data from a single reference laboratory and instrument platform, and the ARUP-derived threshold did not transfer to FlowCAP-II, so multi-institutional pretraining will be needed to establish generalizability. Diagnostic labels were extracted from free-text reports by a large language model and reflect flow-based rather than integrated morphologic and genetic diagnoses, and the cross-site label mappings were imperfect. All evaluations were retrospective, with no comparison against hematopathologists or prospective testing. Because kNN probes depend on neighbor density, they likely understate performance on rare classes relative to task-trained classifiers. The model also lacks direct interpretability. Unlike single-stage attention-based models that highlight predictive cells and markers [14], EventHorizon’s hierarchical architecture complicates event-level attribution, and we are investigating gradient-based attribution and systematic ablation to map predictions to specific cells and markers.

In summary, EventHorizon provides a single, panel-agnostic specimen representation learned from routine clinical flow cytometry without labels. With lightweight classifiers it supports diagnosis, molecular subclassification, and disease-burden estimation, remains robust to assay perturbation, and adapts to external data with minimal calibration. Multi-institutional pretraining, systematic scaling studies, longer event contexts, cell- and marker-level attribution, and prospective evaluation are the next steps toward clinical use.

## Acknowledgements

We thank the ARUP Institute for Research and Innovation for support of this project.

## Data and Code Availability

Model weights and embeddings can be shared upon reasonable request. The pre-training data, including clinical and molecular annotations, are derived from clinical records at ARUP Laboratories and are not publicly available, to protect participant confidentiality. Requests for non-commercial research access may be directed to the corresponding author. Any sharing would require administrative approval from ARUP and the University of Utah and IRB review and a data use agreement, and cannot be guaranteed.

### Use of Generative AI and AI-Assisted Technologies

During the preparation of this manuscript, the authors used Claude (Sonnet 3.5) and Gemini (Flash 3.5) to refine language, improve readability, edit LaTeX markup, and generate Python scripts for data analysis. The authors fully reviewed, verified, and tested all AI-assisted content, code, and analytical outputs, and assume full responsibility for the final manuscript.

## Author Contributions

MMG, MJM, BDO, NCS, and DPN produced the fundamental design of the model. MMG and MJM developed and trained the model. JRJ, RCS, and DPN reviewed flow cytometry data and reports. DPN developed the annotation pipeline. MMG, MJM, and DPN performed data analysis and wrote the manuscript. BDO and DPN supervised the project.

