## Supplementary Information for "EventHorizon: A Foundation Model for Clinical Flow Cytometry"

### Supplementary Information: *EventHorizon: A Foundation Model for Clinical Flow Cytometry*

#### Supplemental Information

This document contains supplementary methods, tables, and figures supporting the main manuscript.

### 1 Supplementary Methods

#### 1.1 Panels and instrumentation

Specimens receive panel combinations driven by specimen type, patient history, and suspected diagnosis. Peripheral blood and bone marrow aspirates always receive B-cell, T-cell, and myeloid screening tubes, and bone marrows additionally receive a plasma cell tube. Other specimen types (e.g., lymph nodes, CSF) receive two general screening tubes (Tissue 1 and 2; Table S3). Based on patient history or the results of initial screening, add-on tubes may be ordered to refine a diagnosis. Panel redundancy across tubes is required to phenotype small populations fully across multiple tubes. All data were acquired on six 3-laser (405, 488, and 533 nm), 10-PMT Beckman Coulter Navios cytometers (Miami, FL). Panels evaluated 9–10 fluorescent markers alongside integrated side scatter and integrated and peak forward scatter intensities.

#### 1.2 Data pre-processing

To prepare our dataset for training our model, the raw clinical data files need preprocessing. We first collect all files output by the cytometers to ingest and standardize/deduplicate metadata such as panel type, fluorescent markers measured, number of events measured, specimen unique IDs, cytometer serial numbers, etc. This allows us to link up multiple tubes/panels from the same specimen and discard unwanted tubes such as duplicates, repeats, failed runs with insufficient measured events, QC tubes, and panels irrelevant for pre-training such as minimum residual disease tests. After selecting this set of "good" data, we run a one-time ingestion to convert and write the raw event and spillover data to disk as safetensors [1].[1] During this one-time ingestion we discard any events for which any detector channel was saturated, since these events cannot be properly compensated. Note, however, that we store events on disk *un*compensated to allow on-the-fly augmentation of spillover/compensation matrices during pre-training, described in the *Training Strategy* section of the main manuscript.

#### 1.3 Diagnostic label extraction

Diagnostic labels were extracted from final flow cytometry reports using Qwen3.5-122B-A10B-FP8 [2] served with vLLM (v0.26.0) [3] on four NVIDIA L40S GPUs. A structured system prompt, supported by 15 few-shot examples, defined 50 multi-label diagnostic categories across 10 disease groups, specimen metadata fields, and uncertainty annotations (Table S2). The schema expands a previously validated BioMedBERT-based system [4] to cover rare diagnostic edge cases and showed high concordance with it on overlapping categories. Hard prompt constraints enforced mutual exclusivity and hierarchical label propagation. Reports were annotated in two stages. First, a deterministic pass (temperature = 0) labeled most reports. Cases failing predefined confidence or consistency checks were escalated to a reasoning pass with a 32 000-token budget, which generated the final annotation alongside an interpretable reasoning trace for manual review. Finally, a deterministic post-processing layer enforced logical consistency rules (e.g., restricting "Normal" to cases in which all disease labels were negative).

#### 1.4 Additional Data Tables

**Table S2**: Annotation labels, prevalence and target definition.

| Label | Prev. (%) | Description |
|---|---|---|
| *Group 1: Normal* | | |
| NORMAL | 30.73 | No flow cytometric abnormality, monoclonality, neoplasm, or dysplasia identified; mutually exclusive with all other labels. |
| *Group 2: B-cell diagnoses* | | |
| CLL | 8.03 | Classic chronic lymphocytic leukemia phenotype (CD5+, CD23+, dim CD20, CD200 bright, light-chain restricted). |
| 5+BNHL | 1.94 | CD5-positive B-cell non-Hodgkin lymphoma not meeting CLL criteria (e.g. mantle cell lymphoma, atypical CLL). |
| 10+BNHL | 3.25 | CD10-positive B-NHL, follicular pattern (follicular lymphoma, Burkitt, other CD10+ B-NHL). |
| DLBCL | 0.30 | Monoclonal B-cell population with any explicit medium/large cell size mention, or report states DLBCL. |

*continued on next page*

[1] This is necessary because reading the raw data from the files output by the cytometers is quite slow, whereas safetensors were explicitly designed for fast reads from disk. While the safetensor format was designed for serializing neural net *model* tensor weights, we find it also works well for efficient storage of tensor *data* on disk if supplemented with structured tables to contain metadata, as long as real-time ingestion of new data is not required as in our use case.

(continued from previous page)

| Label | Prev. (%) | Description |
|---|---|---|
| 5-10-BNHL | 0.65 | Double-negative (CD5− CD10−) B-NHL (marginal zone, lymphoplasmacytic, splenic B-cell lymphomas). |
| B-NHL | 0.71 | Generic/unclassifiable B-cell lymphoma; fallback only when CD5/CD10 status is not assessed. |
| HCL | 0.22 | Classic hairy cell leukemia phenotype (CD11c+, CD25+, CD103+, CD123+, CD200 very bright). |
| HCL_variant | 0.21 | Hairy cell leukemia variant: CD11c+/CD103+ but CD25 and CD123 negative/dim. |
| BALL | 0.67 | B-lymphoblastic leukemia/lymphoma (immature CD19+/CD34+/-/TdT+/- population). |
| 5+MBL | 1.01 | Subthreshold/hedged CD5+ monoclonal B-cell population in peripheral blood (MBL tier of 5+BNHL). |
| 5-10-MBL | 2.24 | Subthreshold/hedged CD5−CD10− monoclonal B-cell population in peripheral blood. |
| MBL | 0.03 | Subthreshold/hedged CD10+ monoclonal B-cell population in peripheral blood. |
| *Group 3: T-cell / NK diagnoses* | | |
| TLPD | 6.97 | T-cell lymphoproliferative disorder; parent label, auto-set when any child label fires. |
| Sezary_CTCL | 0.23 | Sézary syndrome / mycosis fungoides / cutaneous T-cell lymphoma presentation. |
| LGL | 5.65 | Large granular lymphocyte disorder: expanded cytotoxic T/NK-T population with evidence of clonality. |
| TPLL | 0.01 | T-prolymphocytic leukemia (mature T-cell, CD7 bright, CD4+CD8+ co-expression clue). |
| GD_Tcells | 0.71 | Expanded or aberrant gamma-delta T-cell population. |
| EATL | 0.02 | Enteropathy-associated T-cell lymphoma (intraepithelial, CD103+, intestinal context). |
| ALCL | 0.05 | Anaplastic large cell lymphoma: CD30 bright plus explicit large cell size. |
| TALL | 0.15 | T-lymphoblastic leukemia/lymphoma (immature T-cell population); independent of TLPD. |
| NK | 0.40 | Expanded NK-cell population with aberrant phenotype or clonal features. |
| *Group 4: T-cell modifiers* | | |
| TFH | 0.10 | T-follicular helper phenotype (CD4+CD10+PD1+CD200+). |
| T_clonality | 1.33 | T-cell clonality by flow (restricted/skewed TRBC1, or explicit clonality conclusion). |
| DE | 0.15 | Double expressor: CD4+ and CD8+ co-expression on the same T-cell population. |
| CD30_pos | 0.23 | CD30 expression on an abnormal population that does not meet ALCL criteria; always a modifier. |
| *Group 5: Myeloid diagnoses* | | |
| AML | 4.65 | Acute myeloid leukemia: myeloid blasts + equivalents >15% with aberrant myeloid immunophenotype. |
| AUL | 0.03 | Acute undifferentiated leukemia: blasts >15%, fails both myeloid and B/T lineage tests. |
| Myeloid | 0.90 | Abnormal myeloid expansion below the AML blast threshold (≤15%). |
| Monocytic | 3.35 | Monocytic differentiation modifier: AMML component, CMML-type expansion, or reactive monocytosis. |
| MastCell | 0.27 | Abnormal mast cell population (CD117 bright, CD25+) or mastocytosis raised. |
| BPDCN | 0.02 | Blastic plasmacytoid dendritic cell neoplasm (CD123 bright, CD4+, CD56+). |
| PDC_expansion | 0.05 | Increased plasmacytoid dendritic cells insufficient for BPDCN (e.g. CD56-negative or hedged). |
| *Group 6: MPAL* | | |

continued on next page

*(continued from previous page)*

| Label | Prev. (%) | Description |
|---|---|---|
| MPAL_BM | 0.06 | Single blast population co-expressing B-lymphoid and myeloid markers. |
| MPAL_TM | 0.03 | Single blast population co-expressing T-lymphoid and myeloid markers. |
| MPAL_BT | 0.00 | Single blast population co-expressing B-lymphoid and T-lymphoid markers. |
| MPAL_BTM | 0.00 | Single blast population co-expressing B-, T-, and myeloid markers (trilineage). |
| *Group 7: Plasma cell labels* | | |
| PC_neoplasia | 3.14 | Clonal (light-chain restricted) plasma cell population, myeloma-associated phenotype. |
| PC_differentiation | 0.68 | Plasma cell population plasmacytically differentiated from a concurrent B-cell clone. |
| PC_expansion | 0.10 | Polytypic plasma cell expansion above site-specific threshold; no myeloma aberrancy. |
| *Group 8: Other cellular findings* | | |
| Hematogones | 1.38 | Increased benign B-cell precursors with normal, staged maturation (mimics BALL). |
| Thymic | 0.12 | Thymic tissue / hyperplasia (CD1a+, CD4+CD8+ cortical thymocytes, staged maturation). |
| PNH | 0.23 | Paroxysmal nocturnal hemoglobinuria: GPI-anchor deficient population identified. |
| NonHeme | 0.33 | Non-hematologic malignancy suspected (CD45-negative/dim, aberrant CD56/CD200/CD117), plasma cell causes excluded. |
| *Group 9: Clinically notable flags* | | |
| Eos | 2.93 | Eosinophils explicitly reported above 5% of total leukocytes. |
| Left_Shift | 8.09 | Immature myeloid forms in peripheral blood, below AML/Myeloid threshold. |
| High48Ratio | 2.56 | CD4:CD8 ratio $>$ 8:1, read directly from the stated ratio line. |
| Reversed48Ratio | 0.87 | CD4:CD8 ratio $<$ 1:5 (0.2), read directly from the stated ratio line. |
| No20 | 3.21 | No CD20-positive B-cells identified, or CD20 absent where expected. |
| Weak33 | 1.93 | CD33 expression noted as weak/dim/decreased. |
| *Group 10: Quality flags* | | |
| Suboptimal | 32.08 | Any specimen quality concern: viability <80%, high background, limited events, reliability disclaimer. |
| Clotted | 0.57 | Specimen noted as clotted or partially clotted. |
| Hemodilute | 4.20 | Bone marrow specimen noted as hemodilute. |
| Paucicellular | 11.19 | Insufficient cellularity for full analysis. |

| Panel Name | Description | Count |
|---|---|---|
| LL panel B cell | B-NHL screening in PB/BM | 91301 |
| LL panel T cell | T-NHL screening in PB/BM | 86638 |
| LL panel myeloid | Screening for myeloid associated antigens | 86509 |
| LL panel PC | Surface and cytoplasmic markers for plasma cell disorders | 38538 |
| Tissue 1 | B-NHL and BALL screening in Tissues and Fluids | 21499 |
| Tissue 2 | T-NHL screening in Tissues | 21402 |
| Hairy cell mature B add-on | CD5-/CD10- B-NHL evaluation | 8652 |
| AML add-on | Focused evaluation on myeloid blasts | 8446 |
| Mature T Sezary add-on | Additional markers for T-NHL: $TCR$-$\gamma\delta$, CD26, CD30 | 5933 |
| TCC TFH add-on | T cell clonality and T-follicle helper phenotype screening | 3504 |
| Cytoplasmic add-on 1 | Cytoplasmic markers for acute leukemia lineage assignment | 2720 |
| B ALL hematogone add-on | Separate B-ALL from hematogone hyperplasia | 2520 |
| Mega V2 add-on | Megakaryocyte and Erythroid markers | 2513 |
| Mast cell add-on | High sensitivity panel for mast cell disease | 2158 |
| Immature T add-on | Separate T-ALL from thymoma/thymic hyperplasia | 1332 |
| Cytoplasmic add-on 2 | Isotype controls for cytoplasmic add-ons | 1111 |
| B LBL add-on | Additional phenotyping for future B-ALL MRD analysis | 367 |
| Total | | 385143 |

**Table S1**: Summary of our dataset broken down by panel type.

| Panel | FITC | PE | ECD | PC5.5 | PC7 | APC | APC-A700 | APC-A750 | PB | KO |
|---|---|---|---|---|---|---|---|---|---|---|
| AML add-on | CD56 | CD123 | CD5 | CD117 | CD33 | CD7 | CD34 | CD71 | CD15 | CD45 |
| B ALL hematogone add-on | CD58 | CD123 | CD19 | CD22 | CD10 | CRLF2 | CD34 | CD38 | CD20 | CD45 |
| B LBL add-on | CD66b | CD58 | CD19$^{\dagger}$ | CD20 | CD22 | CD24 | CD34 | CD38 | CD10$^{\S}$ | CD45 |
| Cytoplasmic add-on 1 | cTdT | cMPO | CD19$^{\dagger}$ | CD117 | cCD22 | cCD79a | CD34 | cCD3 | empty | CD45 |
| Cytoplasmic add-on 2 | cTdT | cMPO | CD19$^{\dagger}$ | CD117 | empty | cCD79a | CD34 | cCD3 | cCD22$^{\S}$ | CD45 |
| Hairy cell mature B add-on | Kappa$^{*}$ | CD123 | CD19 | CD25 | CD22 | CD103 | CD11c | Lambda$^{\ddagger}$ | CD20 | CD45 |
| Immature T add-on | CD2 | CD1a | CD4 | CD117 | CD33 | CD10 | CD7 | CD3 | CD8 | CD45 |
| LL panel B cell | Kappa | Lambda | CD19 | CD5 | CD23 | CD10 | CD38 | CD200 | CD20 | CD45 |
| LL panel PC | cKappa | cLambda | empty | CD117 | CD56 | CD138 | CD19 | CD38 | CD20 | CD45 |
| LL panel T cell | CD57 | CD7 | CD4 | CD5 | CD2 | CD56 | CD8 | CD3 | CD16 | CD45 |
| LL panel myeloid | CD14 | CD33 | CD13 | CD117 | CD11b | CD38 | CD34 | CD64 | HLA-DR | CD45 |
| Mast cell add-on | empty | CD123 | CD34 | CD25 | CD33 | CD117 | CD30 | CD2 | DAPI | CD45 |
| Mature T Sezary add-on | gamma delta | CD26 | CD4 | CD25 | CD2 | CD7 | CD30 | CD3 | CD8 | CD45 |
| Mega V2 add-on | CD235a | CD41 | CD34 | CD117 | CD61 | CD42b | CD71 | HLA-DR | CD36 | CD45 |
| TCC TFH add-on | CD57 | TRBC1 | CD4 | CD5 | PD-1 | CD2 | CD7 | CD3 | CD200$^{\S}$ | CD8 |
| Tissue 1 | Kappa | Lambda | CD19 | CD5 | CD13 | CD10 | CD34 | CD3 | CD20 | CD45 |
| Tissue 2 | CD2 | CD26 | CD4 | CD5 | CD7 | CD10 | CD30 | CD3 | CD8 | CD45 |

**Table S3**: A complete listing of all antigen/fluorophore combinations for all panels in our dataset. Lowercase "c" prefix indicates a cytoplasmic, rather than surface marker. The cytoplasmic add-on 1/2 were variations of the same panel where cCD22 was moved from BV421 to PE-Cy7. Fluorophore exceptions are: $^{*}$(AlexaFluor488), $^{\dagger}$(PE-CF594), $^{\ddagger}$(APC/Fire 750), $^{\S}$(BV421)

#### 1.5 EventHorizon architecture details

EventHorizon adopts a model architecture featuring a 2-tier hierarchical encoder, similar to FATE [5] but with significant modifications. This strategy unifies the approaches of the flexible cell encoders [6–8] with specimen-level encoders [9–11], keeping the best of both worlds. As shown in Figure 1, EventHorizon's backbone consists of three key elements: a marker tokenizer, a "marker self-attention" Transformer block, and a "cell self-attention" Transformer block.

Our tokenizer uses an embedding lookup table to represent individual marker identities. The scalar expression value for each marker from each cell is concatenated with the appropriate marker embedding, and for each cell, a learnable CLS token is appended to the set of embedded markers for that cell. The CLS token is similar in function to the CLS token used by BERT [12] and Vision Transformers [13]. We use a frozen embedding table with embedding vectors generated similarly to the sin/cos positional encodings from the original Transformer paper [14]. Despite their similarity to standard Transformer positional encoding, these encode only marker identity without positional information. This encoding scheme is a convenient way to generate a table of unique embeddings that are easily distinguishable by Transformers. We experimented with learnable embeddings as used by FATE but found, somewhat to our surprise, that performance was substantially worse than with a frozen table, although this could be worth more thorough experimentation. Channels without a recognized marker identity are retained during compensation and transformation of the full event matrix but are excluded before marker tokenization. This includes both unlabeled open channels and channels explicitly designated as open during marker-name harmonization, so only recognized markers are passed to EventHorizon. We also emphasize that, in contrast with the fixed-panel models, cells from *all panels* for a single specimen can be concatenated together into a single sequence/bag of tokens, since the tokenizer effectively "bakes in" the identities of the measured markers for each cell.

The tokenized representation of the specimen is next passed to a block of "marker self-attention" layers. This is a stack of standard Transformer encoder layers which applies self-attention among markers and CLS token for each cell, but with no attention between different cells (as enforced by a block-diagonal mask). By the end of this block, the CLS token functions as a coarse latent representation of the single cell, having been enriched with information about the cell's measured markers via self-attention. Note that instead of a CLS token, FATE [5] uses cross-attention to pool marker embeddings into a cell-level latent, but in both cases the intent is to produce a cell-level representation in a single shared latent space regardless of what individual markers were measured.

After passing through the stack of marker self-attention layers, the individual marker embeddings have served their purpose and are discarded. Only the cell-level CLS tokens are retained, which now function as cell-level latent representations. Crucially, all cells are mapped into the same shared latent space regardless of what markers were measured on the panel that detected any particular cell. A new *specimen*-level CLS token, learnable and distinct from the cell CLS token used before, is appended to the bag of latent cell tokens, and this bag of cell latents plus specimen CLS is passed through a second stack of standard self-attention Transformer encoder layers, the "cell self-attention" block. This allows *all* the cell latents and specimen CLS token attend to each other, and the latter serves as the final latent representation of the entire specimen after becoming enriched with the full context of all panels measured for the specimen. We expect the output cell latents should also be useful for cell-level downstream tasks, though we do not explore any such use cases in this work.

Optionally, because the marker self-attention layers are far more memory-hungry than the cell self-attention layers, we allow these blocks to have different embedding dimensions to conserve GPU memory in the marker layers while increasing the expressive capacity of the cell layers. A simple linear layer after the marker block but before the cell block projects up from the smaller to the larger latent dimension.

For inference, EventHorizon's complete backbone simply consists of this sequence of marker tokenizer, marker self-attention block, optional up-projection layer, and cell self-attention block, as depicted in Figure 1. The resulting EventHorizon backbone contains 6.5 million trainable parameters; during pre-training, the DINO projection head increases the student model size to 10.9 million trainable parameters.

#### 1.6 EventHorizon pre-training configuration

EventHorizon is pre-trained with a DINO-style teacher–student objective on flow cytometry specimens. Each specimen yields multiple views: differently subsampled events, marker panels, tubes, and augmentations. A shared hierarchical encoder maps each view to a sample embedding, followed by a projection head used only during pre-training. The teacher network is an exponential moving average (EMA) of the student and receives no gradients; its outputs are centered to limit mode collapse. The student is trained to match the teacher via a cross-entropy loss in projection space, plus a weighted kernel density estimator (KDE) regularizer that encourages uniformly spread embeddings. Learning rate, teacher temperature, and KDE weight are linearly warmed or scheduled over training. Augmentations simulate instrument and acquisition variability by perturbing spillover compensation, then applying channel-wise shift and scale transforms; the student receives stronger perturbations than the teacher. This multi-view setup follows the DINO multi-crop idea: a global teacher view is matched against one full student view and multiple smaller cropped student views drawn from the same underlying cells and markers.

A central challenge in adapting DINO-style self-supervised learning to flow cytometry is defining augmentations that are meaningful for cytometric data rather than images. Our pre-training pipeline therefore combines several complementary strategies: flow cytometry-specific transformations adapted from DinoFlow [11], marker masking motivated by prior flow cytometry SSL approaches [5–8, 10], perturbations of the spillover matrix, and a flow cytometry-specific cropping procedure described below.

First, from DinoFlow, we borrow random linear transformations of each marker (i.e., an additive shift and multiplicative scaling) applied to all cells from a single tube. We also aggressively downsample cells to reduce compute cost and increase data diversity; tubes in our dataset typically have anywhere from 30k to 300k events but during training we typically select between 1k and 8k cells per tube. Similar to, but distinct from DinoFlow, we also apply random linear transformations to the spillover matrix, encouraging the model to learn resilience to poorly compensated data.

Second, our masking strategy operates at multiple levels: since each specimen in our dataset always has multiple panels run, for each specimen we can randomly choose some subset of tubes to discard/retain, and for each tube we can randomly choose some number of markers to mask. We mask markers at the tube rather than cell level, and we mask the student views more heavily than the teacher view, encouraging the model to learn meaningful relationships among markers and across tubes/panels. As in DINO, self-distillation is intuitively viewed as distilling the student *from* the target provided by the teacher, hence the teacher should see a more complete view of the specimen than the student.

Our augmentations are conceptually inspired by the multi-crop strategy of SwAV, which presents a model with multiple local views alongside a global image view during training [15]. Translating this idea from images to flow cytometry data is non-trivial. Randomly subsampling cells is analogous to retaining random pixels from an image and does not preserve locality in any meaningful sense. Instead, for each tube we randomly select a marker and retain only cells falling within a randomly chosen quantile interval of that marker, discarding all others. These cropped views contain substantially fewer cells than the corresponding global views, reducing computational cost and enabling multiple independent crops per specimen. In all experiments, the teacher view and at least one student view remain uncropped.

More broadly, our augmentations are not intended to normalize data or eliminate laboratory-specific variation. Rather, they expose the model to realistic sources of technical variability during pre-training, encouraging representations that are robust to such effects. This interpretation aligns with the prevailing view of augmentations in self-supervised learning [16–18]. Recent work has argued that, at sufficiently large scale, explicit invariance-inducing augmentations may be less important than masking alone for preventing overfitting and stabilizing training [19]. Conversely, other studies have demonstrated that self-supervised models do learn meaningful augmentation-induced invariances when such transformations are present during pre-training [20]. Taken together, these findings suggest that augmentation-derived invariances are most beneficial in smaller-data settings and become less critical as dataset scale increases.

This distinction is particularly relevant for flow cytometry. Although our dataset contains more than 50 billion measured cells, it comprises only approximately 100 000 unique specimens, a scale that remains modest compared with contemporary self-supervised training datasets in computer vision and natural language processing. Therefore we suspect that invariance-promoting augmentations remain important for improving optimization, reducing overfitting, and preventing collapse. Conversely, SSL approaches that rely primarily on masking while training on substantially smaller flow cytometry datasets may be disadvantaged [6–8].

As shown in Table S4(a), the encoder is a two-stage transformer: within-cell attention over marker tokens (Stage 1), then across-cell attention to form a sample-level representation (Stage 2). Marker values are concatenated with fixed sinusoidal marker-identity encodings. A two-layer MLP projection head maps the 256-dimensional sample embedding into a 4096-dimensional space where the DINO loss is computed; the projection head is discarded after pre-training.

Table S4(b) summarizes the optimization and DINO parameters. Training uses AdamW with linear warmup, cosine decay to a minimum learning rate, and a batch size of 8 specimens. DINO hyperparameters follow the standard recipe: the student uses a fixed temperature, the teacher temperature increases linearly over 5 000 steps, and teacher weights are updated with momentum 0.994. Teacher outputs are centered with EMA momentum 0.9. The ancillary KDE loss (concentration 1) is added with a weight that ramps from 0.005 to 0.25 over 3 000 steps and is averaged across student views.

Table S4(c) gives the data augmentation parameters. With the specified probabilities, first the spillover matrix is perturbed (additive and multiplicative changes to off-diagonal entries), then events are compensated, then each channel receives an independent random shift and/or scale. Student views use higher application probabilities and larger magnitudes than the teacher, encouraging invariance to mild teacher perturbations and stronger student perturbations. Augmentations are applied before marker subsampling so compensation remains consistent.

(a) Architecture parameters

| Parameter | Value |
|---|---|
| Marker embedding type | Sinusoidal |
| Marker attention heads | 16 |
| Cell attention heads | 16 |
| Number of marker layers | 4 |
| Number of cell layers | 8 |
| Projection head dimension | 4096 |
| Projection head hidden dimension | 1024 |
| Cell embedding dimension | 256 |
| Marker embedding dimension | 64 |
| Marker layer feed-forward dimension | 128 |
| Cell layer feed-forward dimension | 1024 |
| Dropout | 0.1 |

(b) Optimization and DINO parameters

| Parameter | Value |
|---|---|
| Batch size | 8 |
| Maximum learning rate | $5 \times 10^{-5}$ |
| Minimum learning rate | $1 \times 10^{-7}$ |
| Warmup iterations | 2000 |
| Student temperature | 0.15 |
| Teacher momentum | 0.994 |
| Teacher temperature start | 0.04 |
| Teacher temperature end | 0.07 |
| Teacher temperature schedule steps | 5000 |
| Center momentum | 0.9 |
| Ancillary loss type | KDE |
| KDE concentration | 1 |
| KDE loss weight start | 0.005 |
| KDE loss weight end | 0.25 |
| KDE loss weight schedule steps | 3000 |

(c) Data augmentation parameters

| Parameter | Student | Teacher |
|---|---|---|
| Additive spillover perturbation | 0.01 | 0.01 |
| Multiplicative spillover perturbation | 0.10 | 0.05 |
| Scale transform probability | 0.90 | 0.60 |
| Shift transform probability | 0.90 | 0.60 |
| Spillover perturbation probability | 0.90 | 0.60 |
| Scale transform magnitude | 0.20 | 0.10 |
| Shift transform magnitude | 0.30 | 0.10 |

(d) View configuration

| View | Configuration | |
|---|---|---|
| **Teacher** | Number of views | 1 |
| | Events per specimen | 16 384 |
| | Markers per tube | 12 |
| | Tubes | Teacher |
| | Augmentations | Teacher |
| **Student** | Number of views | 1 |
| | Events per specimen | 16 384 |
| | Markers per tube | 10 |
| | Augmentations | Student |
| **Student cropped** | Number of views | 2 |
| | Events per specimen | 1 024 |
| | Markers per tube | 9 |
| | Augmentations | Student |
| | Crop fraction | 0.5 |
| | Crop mode | Interior |
| | Limit markers to | Student view |
| | Reuse cells from | Student view |

**Table S4**: **EventHorizon pre-training configuration.** Summary of architecture, optimization, augmentation, and view-generation parameters used during pre-training.

Table S4(d) gives the views configuration. Each training example defines three view types. The *teacher* view uses all tubes from the specimen, ∼16k events, and 12 markers with mild augmentations. The *student* view independently subsamples three tubes, the same order of events (∼16k), and 10 markers with stronger augmentations. *Student cropped* views are two additional local views of 1 024 events each: events are filtered to an interior quantile window (50% of the range along a randomly chosen marker), then a subset of nine markers is taken from those used in the student view, reusing the same underlying cell indices. The teacher is matched to all student views in the DINO loss; cropped views provide local, multi-crop-style supervision analogous to SwAV and DINOv2 on images [15, 21].

#### 1.7 Model comparison: training, evaluation and hyperparameter-search details

***Data splits.***

All methods share the same specimen IDs and binary targets. The 2019–2025 historical set is partitioned into three folds. The 2026 holdout is used only for final reporting.

***DinoFlow embeddings.***

For DinoFlow [11], we used the frozen backbones pre-trained on the B-cell, T-cell, and myeloid screening panels. Each panel-specific backbone produced a tube-level latent representation, and the three representations were concatenated to form the specimen-level representation used for the triage classification comparison. For the tube-level latent-space analysis, the corresponding panel-specific embeddings were instead analyzed individually.

***Frozen-embedding MLP probes (EventHorizon and DinoFlow).***

Architecture is fixed: embedding → Linear(512)–ReLU–Dropout → Linear(256)–ReLU–Dropout → Linear(1), trained with BCEWithLogitsLoss and class imbalance weight $\text{pos}_{\text{weight}} = n_{\text{neg}}/n_{\text{pos}}$ on the current training rows. A shared hyperparameter configuration (learning rate, dropout, weight decay) is selected once per embedding set by 3-fold CV maximizing macro-mean validation AUPRC over diagnoses. Under the winning configuration, out-of-fold predictions define a single F1-maximizing threshold per label; models are then refit on all historical data for a fixed epoch budget derived from CV best epochs and scored on the 2026 holdout. Paper numbers average three random seeds (mean ± SD).

***Supervised ST-FPS.***

We adapted the model and training code from the original ST-FPS work [22] for specimen-level diagnostic classification. Because the original model produces cell-set representations while our targets are specimen-level, each tube-specific backbone was followed by pooling-by-multi-head-attention (PMA), as in [23]. For the B/T/myeloid screening task, the three pooled tube latents were concatenated and passed to a linear classification head. The model was trained end-to-end from raw B/T/myeloid events with BCEWithLogitsLoss and per-label $\mathrm{pos}_{\mathrm{weight}}$. Shared hyperparameters were selected with a multilabel model on folds $0, 1$ using fold 2 as development (Dev macro AUPRC), then transferred to independent binary models (one per diagnosis) with early stopping on Dev AUPRC and Dev F1 thresholds applied on holdout.

***$k$NN probes.***

As a simpler probe baseline on the same frozen embeddings, we also evaluate distance-weighted cosine $k$NN with shared $k$ chosen by 3-fold CV; holdout results appear in Figure S3.

#### 1.8 Latent-space graph analysis details

For the specimen-level analysis, each node represented one peripheral blood or bone marrow specimen containing the B-cell, T-cell, and myeloid screening tubes. EventHorizon embeddings were generated using all available tubes for each specimen as described in Section 2.4. A $k$-nearest-neighbor graph was constructed from the specimen embeddings using cosine similarity, and graph structure was evaluated using modularity and Leiden community detection [24, 25]. Each specimen was associated with a diagnosis, specimen type, and tube-panel signature, defined as a binary vector indicating which tubes were run. These labels were not used to construct the graph or to identify Leiden communities, but were applied after graph construction to evaluate the organization of the latent space.

To account for the association between tube-panel selection/composition and diagnosis, tube-panel signature modularity was compared with 1000 conditional permutations of tube-panel signature labels occurring more than 50 times, stratified jointly by diagnosis and specimen type [26]. This preserves the major clinical associations between diagnosis, specimen type, and tube-panel selection while breaking any residual association between tube-panel signature and position in the latent graph. Conversely, diagnosis modularity was compared with 1000 conditional permutations of diagnosis labels stratified jointly by tube-panel signature and specimen type. This tests whether diagnosis remains organized in the latent space after accounting for the tubes acquired for each specimen.

Across the dataset, there were 250 unique tube-panel signatures, of which 23 occurred more than 50 times. Leiden communities were evaluated using weighted community purity, defined as the size-weighted fraction of specimens assigned to the most common label within each community, and compared with the corresponding global majority-label baseline. The main analysis used $k = 5$ and Leiden resolution 1.0, with additional parameter settings evaluated in the Supplementary Results.

We also applied this framework to compare the tube-level latent spaces of EventHorizon and DinoFlow using the B-cell, T-cell, and myeloid screening tubes. For this analysis, each tube was embedded separately. EventHorizon was run in single-tube mode, while DinoFlow embeddings were generated using the corresponding panel-specific backbone. Graphs were constructed over the resulting tube embeddings and evaluated with respect to diagnosis, tube type, and cytometer ID using the same modularity and Leiden community-purity measures described above.

### 2 Supplementary Results

#### 2.1 AML Molecular Subclassification

Performance metrics for additional molecular and cytogenetic targets.

**Table S5**: Genotype signal recovered from the flow cytometry case embedding ($2^{17}$ events). Cross-validated (5-fold) logistic regression, embedding → binary label. Cases lacking the relevant assay for a given label are excluded from that label's training and evaluation. AUROC and AUPRC are shown with 95% confidence intervals from 1 000 bootstrap resamples of the out-of-fold predictions. F1, sensitivity, specificity, and precision are reported at each label's own best-F1 threshold from the out-of-fold precision–recall curve. Labels with fewer than 11 positive cases are omitted. Sorted by AUROC.

| Label | Type | AUROC [95% CI] | AUPRC [95% CI] | F1 | Sens. (%) | Spec. (%) | Prec. (%) | $N_+$ | $N_{\text{tested}}$ | Prev. (%) |
|---|---|---|---|---|---|---|---|---|---|---|
| t(15;17) *PML::RARA* | Fusion | 0.991 [0.986–0.996] | 0.810 [0.692–0.902] | 0.745 | 77.8 | 99.0 | 71.4 | 45 | 1473 | 3.1 |
| t(8;21) *RUNX1::RUNX1T1* | Fusion | 0.981 [0.961–0.996] | 0.613 [0.350–0.832] | 0.600 | 69.2 | 99.1 | 52.9 | 13 | 912 | 1.4 |
| *DDX41* | Gene | 0.970 [0.954–0.982] | 0.537 [0.399–0.668] | 0.571 | 68.0 | 98.2 | 49.3 | 50 | 2013 | 2.5 |
| inv(16)/t(16;16) *CBFB::MYH11* | Fusion | 0.967 [0.914–0.996] | 0.674 [0.480–0.875] | 0.739 | 77.3 | 99.2 | 70.8 | 22 | 912 | 2.4 |
| *CALR* | Gene | 0.893 [0.833–0.940] | 0.285 [0.176–0.436] | 0.342 | 59.1 | 95.8 | 24.1 | 44 | 2015 | 2.2 |
| *NPM1* mutated, no other cytogenetic abn. | Composite | 0.887 [0.857–0.917] | 0.525 [0.444–0.609] | 0.562 | 69.9 | 92.9 | 47.0 | 166 | 2013 | 8.2 |
| *NPM1* | Gene | 0.869 [0.836–0.901] | 0.519 [0.439–0.602] | 0.539 | 59.5 | 94.2 | 49.3 | 173 | 2014 | 8.6 |
| *KIT* | Gene | 0.863 [0.808–0.912] | 0.195 [0.125–0.310] | 0.295 | 45.1 | 95.8 | 21.9 | 51 | 2014 | 2.5 |
| *BRAF* | Gene | 0.861 [0.775–0.940] | 0.246 [0.084–0.452] | 0.320 | 21.1 | 99.9 | 66.7 | 19 | 2015 | 0.9 |
| *KMT2A* rearranged | Fusion | 0.843 [0.769–0.900] | 0.317 [0.209–0.467] | 0.429 | 51.1 | 95.3 | 36.9 | 47 | 912 | 5.2 |
| *LUC7L2* | Gene | 0.841 [0.710–0.947] | 0.229 [0.053–0.460] | 0.320 | 23.5 | 99.8 | 50.0 | 17 | 2013 | 0.8 |
| *NUP98* rearranged | Fusion | 0.828 [0.637–0.966] | 0.267 [0.059–0.529] | 0.293 | 50.0 | 97.4 | 20.7 | 12 | 912 | 1.3 |
| *STAG2* | Gene | 0.827 [0.791–0.863] | 0.396 [0.326–0.482] | 0.467 | 56.8 | 92.2 | 39.7 | 169 | 2030 | 8.3 |
| *CEBPA* | Gene | 0.815 [0.775–0.852] | 0.190 [0.139–0.275] | 0.276 | 27.2 | 96.7 | 28.1 | 92 | 2019 | 4.6 |
| *SMC1A* | Gene | 0.812 [0.710–0.898] | 0.089 [0.032–0.239] | 0.214 | 31.6 | 98.4 | 16.2 | 19 | 2012 | 0.9 |
| del(17p) | CNV | 0.810 [0.684–0.908] | 0.210 [0.080–0.409] | 0.269 | 64.3 | 85.4 | 17.0 | 14 | 315 | 4.4 |
| *FLT3* | Gene | 0.808 [0.783–0.831] | 0.427 [0.375–0.488] | 0.493 | 70.2 | 79.1 | 38.0 | 312 | 2028 | 15.4 |
| *ASXL1* | Gene | 0.806 [0.784–0.828] | 0.581 [0.535–0.630] | 0.598 | 69.0 | 80.2 | 52.8 | 503 | 2066 | 24.3 |
| *MECOM* (EVI1) rearranged | Fusion | 0.804 [0.701–0.899] | 0.167 [0.069–0.323] | 0.246 | 34.8 | 96.2 | 19.0 | 23 | 912 | 2.5 |
| *ETNK1* | Gene | 0.800 [0.708–0.883] | 0.099 [0.033–0.208] | 0.132 | 37.5 | 94.8 | 8.0 | 24 | 2011 | 1.2 |
| complex karyotype (3+ copy-number abn.) | Composite | 0.797 [0.754–0.836] | 0.327 [0.261–0.415] | 0.452 | 54.7 | 87.2 | 38.6 | 117 | 916 | 12.8 |
| del(5q) / monosomy 5 | CNV | 0.793 [0.752–0.831] | 0.452 [0.374–0.539] | 0.508 | 62.9 | 83.3 | 42.6 | 151 | 917 | 16.5 |
| *FLT3*-ITD | Gene | 0.792 [0.757–0.824] | 0.352 [0.296–0.417] | 0.438 | 53.4 | 88.8 | 37.1 | 223 | 2023 | 11.0 |
| *GATA2* | Gene | 0.787 [0.742–0.827] | 0.186 [0.135–0.267] | 0.277 | 36.1 | 93.7 | 22.4 | 97 | 2019 | 4.8 |
| *ZRSR2* | Gene | 0.779 [0.701–0.844] | 0.118 [0.061–0.229] | 0.182 | 11.9 | 99.6 | 38.5 | 42 | 2019 | 2.1 |
| *BCOR* | Gene | 0.775 [0.729–0.820] | 0.181 [0.136–0.249] | 0.289 | 39.3 | 92.2 | 22.8 | 112 | 2020 | 5.5 |
| *TP53* | Gene | 0.775 [0.749–0.801] | 0.472 [0.423–0.530] | 0.507 | 61.4 | 80.4 | 43.2 | 399 | 2041 | 19.5 |
| other FISH abnormality | CNV | 0.774 [0.692–0.842] | 0.145 [0.094–0.236] | 0.268 | 38.6 | 92.4 | 20.5 | 44 | 912 | 4.8 |
| *SRSF2* | Gene | 0.766 [0.731–0.798] | 0.379 [0.318–0.443] | 0.425 | 47.9 | 88.5 | 38.2 | 263 | 2034 | 12.9 |
| MDS-related genes (any of 10) | Composite | 0.764 [0.743–0.785] | 0.728 [0.698–0.758] | 0.706 | 83.0 | 54.6 | 61.5 | 979 | 2102 | 46.6 |
| del(7q) / monosomy 7 | CNV | 0.763 [0.713–0.809] | 0.335 [0.267–0.432] | 0.427 | 49.6 | 88.2 | 37.5 | 115 | 917 | 12.5 |
| *RUNX1* | Gene | 0.761 [0.733–0.788] | 0.410 [0.367–0.465] | 0.475 | 66.4 | 74.9 | 36.9 | 372 | 2052 | 18.1 |
| *SF3B1* | Gene | 0.755 [0.698–0.808] | 0.152 [0.101–0.225] | 0.246 | 27.7 | 95.8 | 22.1 | 83 | 2018 | 4.1 |
| *SETBP1* | Gene | 0.755 [0.685–0.822] | 0.178 [0.099–0.279] | 0.249 | 34.9 | 95.3 | 19.3 | 63 | 2021 | 3.1 |
| *SH2B3* | Gene | 0.755 [0.642–0.859] | 0.063 [0.021–0.183] | 0.154 | 15.0 | 99.2 | 15.8 | 20 | 2012 | 1.0 |
| *NOTCH1* | Gene | 0.753 [0.558–0.930] | 0.299 [0.131–0.594] | 0.500 | 38.9 | 99.8 | 70.0 | 18 | 2014 | 0.9 |
| *CBL* | Gene | 0.752 [0.686–0.812] | 0.126 [0.076–0.205] | 0.223 | 37.5 | 93.5 | 15.9 | 64 | 2021 | 3.2 |
| *IDH2* | Gene | 0.750 [0.712–0.788] | 0.272 [0.217–0.338] | 0.336 | 41.2 | 89.7 | 28.4 | 182 | 2025 | 9.0 |
| *PHF6* | Gene | 0.742 [0.695–0.791] | 0.149 [0.116–0.205] | 0.247 | 39.6 | 89.5 | 18.0 | 111 | 2020 | 5.5 |
| trisomy 7 / chr7 gain | CNV | 0.738 [0.576–0.890] | 0.215 [0.036–0.464] | 0.316 | 23.1 | 99.7 | 50.0 | 13 | 912 | 1.4 |
| *JAK2* | Gene | 0.732 [0.677–0.786] | 0.252 [0.177–0.350] | 0.298 | 26.2 | 97.2 | 34.6 | 107 | 2023 | 5.3 |
| *FLT3*-TKD | Gene | 0.719 [0.651–0.781] | 0.096 [0.057–0.152] | 0.151 | 42.0 | 85.3 | 9.2 | 69 | 2014 | 3.4 |
| *WT1* | Gene | 0.718 [0.675–0.762] | 0.199 [0.155–0.264] | 0.273 | 25.2 | 95.1 | 29.8 | 155 | 2022 | 7.7 |
| *CUX1* | Gene | 0.713 [0.628–0.782] | 0.101 [0.057–0.196] | 0.194 | 16.7 | 98.5 | 23.1 | 54 | 2015 | 2.7 |
| *CSF3R* | Gene | 0.707 [0.589–0.819] | 0.037 [0.017–0.102] | 0.113 | 19.0 | 97.7 | 8.0 | 21 | 2012 | 1.0 |
| *NRAS* | Gene | 0.700 [0.659–0.736] | 0.241 [0.201–0.298] | 0.319 | 52.6 | 77.2 | 22.9 | 230 | 2020 | 11.4 |
| *TET2* | Gene | 0.698 [0.668–0.725] | 0.390 [0.344–0.444] | 0.418 | 53.8 | 74.9 | 34.2 | 400 | 2047 | 19.5 |
| *EZH2* | Gene | 0.692 [0.637–0.747] | 0.180 [0.122–0.254] | 0.242 | 31.9 | 92.2 | 19.5 | 113 | 2030 | 5.6 |
| *U2AF1* | Gene | 0.691 [0.647–0.732] | 0.201 [0.156–0.265] | 0.279 | 34.6 | 90.1 | 23.3 | 162 | 2026 | 8.0 |
| *IDH1* | Gene | 0.687 [0.629–0.739] | 0.114 [0.085–0.161] | 0.212 | 38.9 | 87.1 | 14.5 | 108 | 2018 | 5.4 |
| *KDM6A* | Gene | 0.685 [0.532–0.831] | 0.092 [0.027–0.245] | 0.229 | 21.1 | 99.4 | 25.0 | 19 | 2014 | 0.9 |
| *BCORL1* | Gene | 0.684 [0.590–0.774] | 0.089 [0.039–0.182] | 0.168 | 28.6 | 95.5 | 11.9 | 42 | 2015 | 2.1 |
| *PPM1D* | Gene | 0.683 [0.581–0.775] | 0.069 [0.032–0.163] | 0.126 | 17.9 | 96.7 | 9.7 | 39 | 2015 | 1.9 |
| chr5 gain | CNV | 0.671 [0.492–0.828] | 0.037 [0.014–0.114] | 0.109 | 25.0 | 95.6 | 7.0 | 12 | 912 | 1.3 |
| *MPL* | Gene | 0.668 [0.551–0.779] | 0.043 [0.020–0.114] | 0.115 | 17.2 | 97.3 | 8.6 | 29 | 2013 | 1.4 |
| *ASXL2* | Gene | 0.667 [0.562–0.770] | 0.052 [0.023–0.138] | 0.098 | 6.2 | 99.6 | 22.2 | 32 | 2016 | 1.6 |
| *PRPF8* | Gene | 0.666 [0.547–0.772] | 0.018 [0.007–0.052] | 0.063 | 12.5 | 97.7 | 4.3 | 16 | 2014 | 0.8 |
| *ETV6* | Gene | 0.658 [0.567–0.735] | 0.050 [0.029–0.094] | 0.128 | 14.0 | 97.7 | 11.8 | 43 | 2015 | 2.1 |
| *DNMT3A* | Gene | 0.648 [0.615–0.679] | 0.295 [0.258–0.340] | 0.368 | 64.1 | 61.5 | 25.8 | 351 | 2036 | 17.2 |
| *KRAS* | Gene | 0.642 [0.591–0.691] | 0.108 [0.078–0.159] | 0.158 | 27.4 | 86.6 | 11.1 | 117 | 2023 | 5.8 |
| *RAD21* | Gene | 0.634 [0.520–0.733] | 0.036 [0.014–0.127] | 0.074 | 7.7 | 98.7 | 7.1 | 26 | 2013 | 1.3 |
| *PTPN11* | Gene | 0.620 [0.566–0.674] | 0.078 [0.055–0.127] | 0.132 | 11.0 | 97.4 | 16.7 | 91 | 2017 | 4.5 |
| *SMC3* | Gene | 0.617 [0.457–0.781] | 0.010 [0.005–0.024] | 0.029 | 50.0 | 80.4 | 1.5 | 12 | 2013 | 0.6 |
| copy-neutral LOH | CNV | 0.614 [0.525–0.699] | 0.250 [0.183–0.358] | 0.377 | 60.0 | 66.5 | 27.5 | 55 | 315 | 17.5 |
| trisomy 8 | CNV | 0.441 [0.303–0.571] | 0.047 [0.027–0.091] | 0.108 | 68.8 | 40.8 | 5.9 | 16 | 315 | 5.1 |

#### 2.2 Spectral flow cytometry evaluation

We evaluated spectral flow cytometry files acquired from 28 residual clinical specimens using an unvalidated experimental 37-color panel on a Cytek Aurora cytometer. This panel lacks several add-on markers from our conventional panels and includes an un-pretrained viability dye (Viakrome 808), which was excluded prior to tokenization so as not to affect downstream embeddings. The same 28 specimens were concurrently run on conventional 10-color Navios panels and incorporated into both pre-training and classifier training.

A blinded hematopathologist (JRJ) assigned free-text diagnoses to each spectral specimen (Figure S1), which were mapped to standardized ARUP labels to establish the evaluation benchmark (e.g., "AMML with abn pDC" to AML, Monocytic, and PDC_expansion; "No evidence of disease" to NORMAL). Correct labels are outlined in red in Figure S1.

Spectral embeddings were generated following Section 2.4. To prevent data leakage, each specimen's matched Navios embedding was excluded from the 117 697 quality-filtered reference cases. We retrieved the 25 nearest neighbors by cosine similarity and aggregated reference labels via similarity-weighted voting. Predictions were evaluated at multiple retrieval depths (top-voted, top-3, single nearest, or any within 25 nearest neighbors).

Overall, top-voted agreement reached 46% (13/28), top-3 coverage reached 61% (17/28), top-neighbor coverage reached 71% (20/28), and 1-NN accuracy reached 29% (8/28). By disease category, top-voted accuracy was:

- 6 of 7 myeloid and monocytic neoplasms,
- 1 of 2 acute leukemias with B/myeloid phenotype,
- 2 of 3 plasma cell neoplasms,
- 3 of 4 T- and NK-cell populations,
- 0 of 5 B-cell neoplasms and other specimens, and
- 1 of 7 cases with no evidence of disease.

Underperformance in B-cell neoplasms largely reflects panel limitations: key diagnostic markers like CD103 were omitted, and poor reagent performance for $\kappa$ and $\lambda$ light chains obscured clonality assessment. Per-dimension moment-matching domain adaptation yielded no material change (14/28 top-voted matches; 22/28 within 25 nearest neighbors). Figure S1 details the full 25-neighbor breakdown for each specimen.

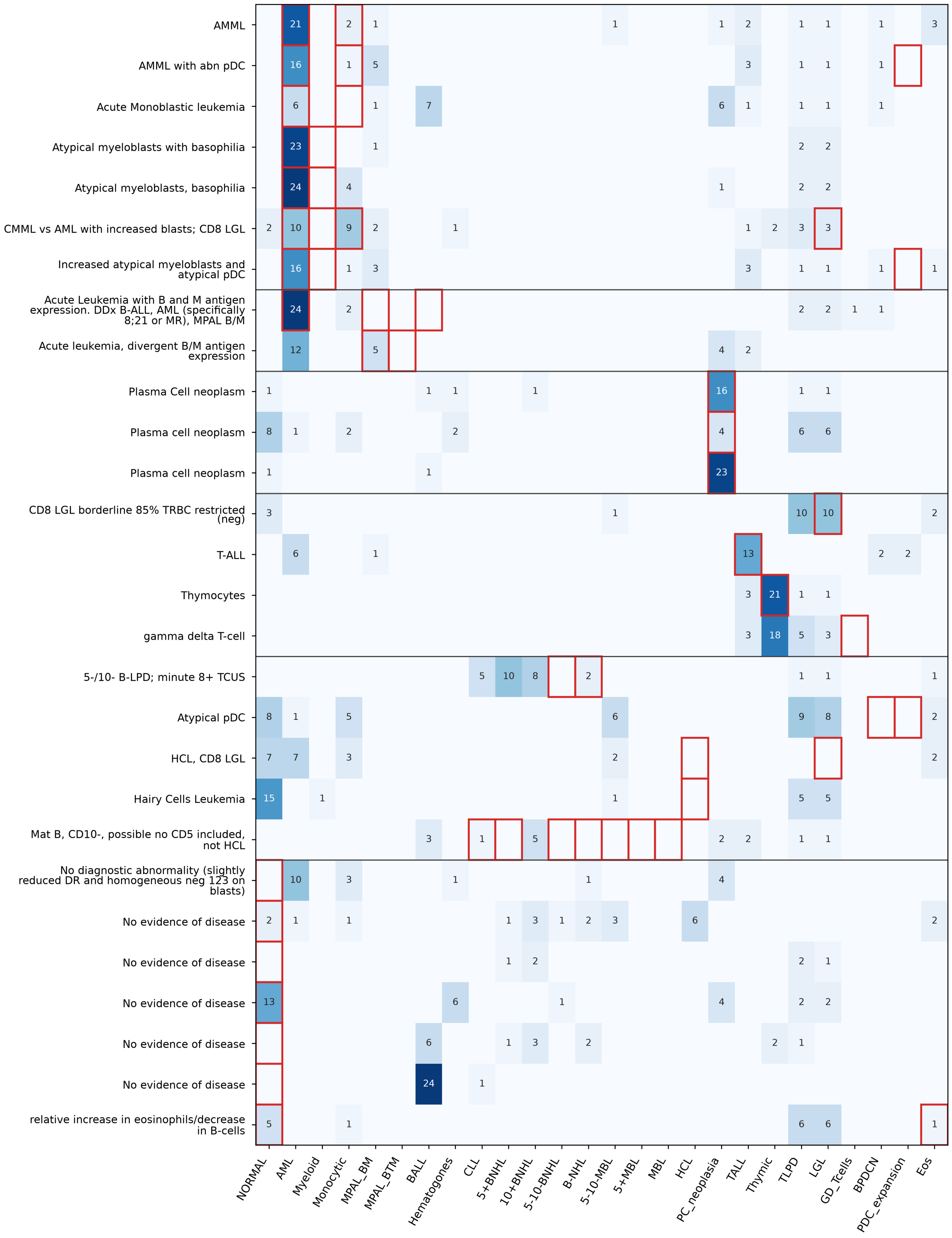

**Fig. S1**: **Labels among the 25 nearest ARUP neighbors of each spectral flow specimen.** Rows correspond to the 28 specimens, labeled with the blinded hematopathologist's (JRJ) full comment and grouped by diagnostic category. Columns denote ARUP diagnostic labels. Each cell indicates the number of 25 nearest neighbors (cosine similarity, excluding matched Navios runs) carrying that label. Red boxes indicate labels consistent with the reviewer's diagnosis. Labels with fewer than four occurrences across all specimens are omitted unless expected.

#### 2.3 Temporal data split performance - full data

Full per-label performance for the 2026 validation cohort is shown below for all five classifiers. Results are reported for the 28 diagnostic labels with at least 10 positive cases, including AUPRC and $F_1$ with bootstrap 95% confidence intervals. The highest AUPRC and $F_1$ for each label are bolded independently.

**Table S6**: Full per-label 2026 validation-set performance on 10 901 cases (AUPRC and F1, both with bootstrap 95% CI, 1 000 resamples) for all five classifiers, labels with $\geq 10$ positives out of $n = 10\,901$, sorted by prevalence. The highest AUPRC and F1 for each label are bolded independently,

| Label | $n_+$ | Model | AUPRC [95% CI] | F1 [95% CI] |
|---|---|---|---|---|
| NORMAL | 3270 | kNN | 0.799 [0.784–0.813] | 0.745 [0.734–0.756] |
| | | LogReg | 0.814 [0.799–0.828] | 0.752 [0.742–0.765] |
| | | RF | 0.788 [0.775–0.802] | 0.727 [0.716–0.739] |
| | | XGB | 0.817 [0.803–0.832] | 0.755 [0.745–0.766] |
| | | MLP | **0.830 [0.817–0.844]** | **0.764 [0.753–0.774]** |
| CLL | 865 | kNN | 0.953 [0.941–0.964] | 0.880 [0.865–0.896] |
| | | LogReg | 0.970 [0.962–0.978] | 0.910 [0.896–0.924] |
| | | RF | 0.958 [0.949–0.967] | 0.884 [0.869–0.899] |
| | | XGB | 0.975 [0.967–0.981] | 0.913 [0.899–0.926] |
| | | MLP | **0.978 [0.972–0.984]** | **0.925 [0.911–0.937]** |
| AML | 474 | kNN | 0.936 [0.916–0.953] | 0.868 [0.843–0.890] |
| | | LogReg | 0.937 [0.917–0.955] | 0.875 [0.852–0.896] |
| | | RF | 0.934 [0.912–0.953] | 0.878 [0.855–0.900] |
| | | XGB | 0.941 [0.924–0.956] | 0.873 [0.851–0.892] |
| | | MLP | **0.944 [0.926–0.960]** | **0.883 [0.861–0.905]** |
| TLPD | 435 | kNN | 0.610 [0.560–0.662] | 0.549 [0.515–0.581] |
| | | LogReg | 0.607 [0.560–0.656] | 0.540 [0.508–0.571] |
| | | RF | 0.611 [0.562–0.660] | 0.496 [0.466–0.528] |
| | | XGB | 0.619 [0.572–0.662] | 0.563 [0.530–0.595] |
| | | MLP | **0.641 [0.596–0.685]** | **0.585 [0.550–0.616]** |
| Monocytic | 420 | kNN | 0.596 [0.551–0.637] | 0.532 [0.485–0.582] |
| | | LogReg | 0.680 [0.637–0.722] | **0.645 [0.607–0.681]** |
| | | RF | 0.612 [0.569–0.657] | 0.554 [0.510–0.595] |
| | | XGB | 0.674 [0.631–0.719] | 0.620 [0.579–0.663] |
| | | MLP | **0.694 [0.650–0.737]** | 0.635 [0.596–0.669] |
| 5-10-MBL | 348 | kNN | 0.419 [0.364–0.475] | 0.510 [0.466–0.553] |
| | | LogReg | 0.525 [0.471–0.586] | 0.462 [0.414–0.507] |
| | | RF | 0.485 [0.430–0.544] | 0.525 [0.480–0.566] |
| | | XGB | 0.505 [0.452–0.564] | 0.510 [0.465–0.554] |
| | | MLP | **0.583 [0.530–0.641]** | **0.594 [0.556–0.632]** |
| LGL | 299 | kNN | 0.511 [0.455–0.574] | 0.446 [0.409–0.484] |
| | | LogReg | 0.509 [0.447–0.566] | **0.480 [0.439–0.520]** |
| | | RF | **0.535 [0.478–0.601]** | 0.418 [0.386–0.451] |
| | | XGB | 0.490 [0.431–0.558] | 0.439 [0.400–0.477] |
| | | MLP | 0.507 [0.448–0.572] | 0.477 [0.437–0.517] |
| T_clonality | 283 | kNN | 0.437 [0.381–0.496] | **0.583 [0.538–0.623]** |
| | | LogReg | 0.494 [0.437–0.560] | 0.567 [0.522–0.609] |
| | | RF | 0.456 [0.401–0.516] | 0.543 [0.498–0.587] |
| | | XGB | 0.499 [0.443–0.566] | 0.527 [0.481–0.570] |
| | | MLP | **0.504 [0.450–0.570]** | 0.546 [0.502–0.591] |
| PC_neoplasia | 262 | kNN | 0.625 [0.565–0.690] | 0.605 [0.557–0.650] |
| | | LogReg | 0.720 [0.664–0.776] | 0.712 [0.665–0.756] |
| | | RF | 0.630 [0.568–0.694] | 0.649 [0.597–0.696] |
| | | XGB | 0.666 [0.601–0.729] | 0.668 [0.616–0.716] |
| | | MLP | **0.770 [0.726–0.816]** | **0.718 [0.674–0.763]** |
| 5+BNHL | 245 | kNN | 0.426 [0.366–0.498] | 0.417 [0.355–0.477] |
| | | LogReg | 0.550 [0.487–0.617] | 0.531 [0.478–0.580] |
| | | RF | 0.494 [0.431–0.555] | 0.486 [0.429–0.535] |
| | | XGB | 0.537 [0.470–0.603] | 0.519 [0.463–0.567] |

Table S6 – continued

| Label | $n_+$ | Model | AUPRC [95% CI] | F1 [95% CI] |
|---|---|---|---|---|
| | | MLP | **0.642 [0.583–0.702]** | **0.605 [0.552–0.657]** |
| Eos | 229 | kNN | 0.213 [0.161–0.274] | 0.284 [0.236–0.338] |
| | | LogReg | 0.598 [0.532–0.661] | **0.573 [0.526–0.618]** |
| | | RF | 0.235 [0.189–0.305] | 0.314 [0.267–0.363] |
| | | XGB | 0.465 [0.399–0.526] | 0.458 [0.406–0.512] |
| | | MLP | **0.607 [0.548–0.669]** | 0.552 [0.498–0.604] |
| 10+BNHL | 197 | kNN | 0.505 [0.432–0.587] | 0.571 [0.515–0.622] |
| | | LogReg | 0.587 [0.514–0.664] | 0.589 [0.539–0.637] |
| | | RF | 0.523 [0.454–0.607] | 0.528 [0.479–0.579] |
| | | XGB | 0.603 [0.535–0.678] | **0.600 [0.549–0.647]** |
| | | MLP | **0.663 [0.592–0.736]** | 0.590 [0.541–0.635] |
| 5-10-BNHL | 116 | kNN | 0.055 [0.034–0.094] | 0.044 [0.000–0.095] |
| | | LogReg | **0.152 [0.114–0.212]** | 0.164 [0.091–0.237] |
| | | RF | 0.124 [0.086–0.190] | **0.215 [0.163–0.267]** |
| | | XGB | 0.141 [0.101–0.204] | 0.086 [0.028–0.155] |
| | | MLP | 0.127 [0.092–0.182] | 0.068 [0.015–0.127] |
| 5+MBL | 85 | kNN | 0.286 [0.217–0.389] | 0.389 [0.316–0.456] |
| | | LogReg | 0.330 [0.252–0.430] | 0.364 [0.289–0.436] |
| | | RF | 0.331 [0.261–0.434] | 0.413 [0.337–0.482] |
| | | XGB | 0.301 [0.233–0.398] | 0.402 [0.313–0.481] |
| | | MLP | **0.360 [0.270–0.465]** | **0.438 [0.363–0.509]** |
| GD_Tcells | 75 | kNN | 0.131 [0.079–0.209] | 0.162 [0.098–0.225] |
| | | LogReg | 0.113 [0.085–0.155] | **0.207 [0.146–0.265]** |
| | | RF | **0.168 [0.105–0.256]** | 0.182 [0.129–0.240] |
| | | XGB | 0.132 [0.093–0.192] | 0.191 [0.144–0.239] |
| | | MLP | 0.152 [0.105–0.222] | 0.205 [0.146–0.264] |
| PC_differentiation | 67 | kNN | 0.176 [0.107–0.274] | 0.236 [0.131–0.333] |
| | | LogReg | **0.253 [0.176–0.373]** | **0.351 [0.248–0.444]** |
| | | RF | 0.214 [0.143–0.319] | 0.265 [0.157–0.368] |
| | | XGB | 0.236 [0.155–0.339] | 0.212 [0.112–0.315] |
| | | MLP | 0.245 [0.163–0.362] | 0.309 [0.217–0.393] |
| DLBCL | 60 | kNN | 0.241 [0.149–0.366] | 0.370 [0.257–0.483] |
| | | LogReg | 0.208 [0.142–0.319] | 0.241 [0.130–0.349] |
| | | RF | **0.348 [0.225–0.481]** | **0.382 [0.258–0.505]** |
| | | XGB | 0.280 [0.186–0.404] | 0.253 [0.124–0.377] |
| | | MLP | 0.315 [0.213–0.450] | 0.231 [0.110–0.357] |
| Myeloid | 50 | kNN | 0.339 [0.207–0.500] | 0.382 [0.257–0.488] |
| | | LogReg | 0.208 [0.121–0.340] | 0.315 [0.200–0.423] |
| | | RF | **0.383 [0.257–0.527]** | **0.460 [0.319–0.571]** |
| | | XGB | 0.298 [0.173–0.441] | 0.378 [0.250–0.495] |
| | | MLP | 0.331 [0.210–0.484] | 0.387 [0.269–0.488] |
| BALL | 43 | kNN | 0.948 [0.885–0.994] | 0.899 [0.828–0.957] |
| | | LogReg | 0.935 [0.862–0.988] | **0.918 [0.841–0.972]** |
| | | RF | **0.966 [0.922–0.996]** | 0.899 [0.828–0.961] |
| | | XGB | 0.953 [0.899–0.994] | 0.894 [0.815–0.958] |
| | | MLP | 0.957 [0.904–0.996] | 0.886 [0.812–0.946] |
| B-NHL | 31 | kNN | 0.051 [0.024–0.120] | 0.088 [0.029–0.158] |
| | | LogReg | **0.075 [0.041–0.158]** | 0.108 [0.054–0.167] |
| | | RF | 0.057 [0.033–0.107] | **0.124 [0.055–0.199]** |
| | | XGB | 0.061 [0.037–0.112] | 0.093 [0.032–0.164] |
| | | MLP | 0.064 [0.033–0.140] | 0.110 [0.034–0.189] |
| NonHeme | 30 | kNN | 0.276 [0.166–0.463] | 0.381 [0.231–0.523] |
| | | LogReg | 0.264 [0.153–0.436] | 0.322 [0.187–0.442] |
| | | RF | 0.330 [0.192–0.525] | **0.476 [0.321–0.620]** |
| | | XGB | **0.405 [0.238–0.595]** | 0.447 [0.300–0.578] |
| | | MLP | 0.311 [0.178–0.486] | 0.448 [0.286–0.595] |
| Hematogones | 30 | kNN | 0.201 [0.105–0.359] | 0.209 [0.123–0.289] |
| | | LogReg | 0.200 [0.095–0.373] | 0.238 [0.143–0.336] |

Table S6 – continued

| Label | $n_+$ | Model | AUPRC [95% CI] | F1 [95% CI] |
|---|---|---|---|---|
| | | RF | 0.213 [0.096–0.374] | 0.182 [0.105–0.264] |
| | | XGB | 0.169 [0.086–0.329] | 0.204 [0.115–0.286] |
| | | MLP | **0.255 [0.134–0.417]** | **0.256 [0.151–0.357]** |
| MastCell | 25 | kNN | 0.196 [0.072–0.369] | 0.222 [0.053–0.391] |
| | | LogReg | 0.189 [0.110–0.356] | 0.250 [0.087–0.409] |
| | | RF | 0.197 [0.091–0.384] | 0.286 [0.128–0.425] |
| | | XGB | 0.292 [0.139–0.493] | **0.400 [0.167–0.593]** |
| | | MLP | **0.324 [0.183–0.549]** | 0.316 [0.111–0.500] |
| HCL | 22 | kNN | 0.496 [0.300–0.724] | 0.296 [0.087–0.522] |
| | | LogReg | 0.702 [0.474–0.862] | 0.652 [0.476–0.792] |
| | | RF | 0.429 [0.208–0.640] | 0.400 [0.167–0.606] |
| | | XGB | 0.705 [0.526–0.871] | **0.683 [0.500–0.833]** |
| | | MLP | **0.784 [0.608–0.917]** | 0.667 [0.454–0.829] |
| NK | 20 | kNN | 0.309 [0.125–0.518] | 0.364 [0.148–0.564] |
| | | LogReg | 0.236 [0.084–0.459] | 0.342 [0.138–0.511] |
| | | RF | 0.354 [0.161–0.590] | **0.385 [0.189–0.548]** |
| | | XGB | 0.342 [0.148–0.564] | 0.353 [0.174–0.512] |
| | | MLP | **0.450 [0.233–0.665]** | 0.378 [0.174–0.565] |
| HCL_variant | 19 | kNN | 0.124 [0.049–0.269] | **0.274 [0.133–0.400]** |
| | | LogReg | 0.076 [0.036–0.174] | 0.080 [0.000–0.186] |
| | | RF | **0.172 [0.059–0.340]** | 0.130 [0.064–0.197] |
| | | XGB | 0.131 [0.049–0.331] | 0.136 [0.061–0.211] |
| | | MLP | 0.129 [0.059–0.281] | 0.234 [0.130–0.346] |
| CD30_pos | 16 | kNN | 0.120 [0.046–0.313] | 0.174 [0.000–0.364] |
| | | LogReg | 0.108 [0.023–0.273] | 0.093 [0.000–0.222] |
| | | RF | 0.176 [0.061–0.424] | 0.269 [0.095–0.425] |
| | | XGB | 0.127 [0.046–0.312] | 0.154 [0.000–0.348] |
| | | MLP | **0.219 [0.070–0.424]** | **0.286 [0.080–0.462]** |
| TFH | 14 | kNN | 0.009 [0.001–0.043] | 0.000 [0.000–0.000] |
| | | LogReg | 0.070 [0.009–0.262] | **0.125 [0.000–0.286]** |
| | | RF | 0.023 [0.003–0.099] | 0.000 [0.000–0.000] |
| | | XGB | **0.089 [0.008–0.260]** | **0.125 [0.000–0.364]** |
| | | MLP | 0.049 [0.020–0.114] | 0.065 [0.000–0.188] |

#### 2.4 Comparison with other models

The main triage evaluation included 33 diagnostic labels with at least 10 positive cases in the 2026 holdout cohort. Figure S2 shows the full per-label results for EventHorizon, DinoFlow, and ST-FPS using the primary MLP-probe comparison. EventHorizon is shown using either the three triage tubes alone or all tubes retained for each specimen after preprocessing. To assess whether the relative performance of the frozen representations depended on probe complexity, we additionally evaluated distance-weighted cosine $k$-nearest-neighbor probes on the EventHorizon and DinoFlow embeddings. Figure S3 shows the corresponding full per-label results using 5-NN probes.

The per-label MLP and $k$NN results treat each diagnosis independently and therefore do not show which labels are confused with one another. To provide that view, we restricted the 2026 holdout to specimens with exactly one gold label among the eight diagnoses used in the main-text comparison (NORMAL, CLL, AML, BALL, Myeloid, LGL, 5-10-BNHL, and plasma-cell neoplasia; $n = 4\,729$). For each specimen we scored the eight corresponding binary models, assigned a single predicted class by $\arg\max$ of those probabilities, and computed row-normalized confusion matrices (Figure S4). This construction uses the same trained heads and seeds as the bar-plot comparison (it is not a separately trained multiclass model), and multi-label specimens are excluded. Frequent exclusive diagnoses (NORMAL, CLL, AML, BALL) are recovered with high row accuracy for all methods. Errors on rarer exclusive labels more often fall into NORMAL, particularly for ST-FPS under $\arg\max$, which is expected when the highest-scoring head is used in place of the per-label F1 thresholds from the primary evaluation.

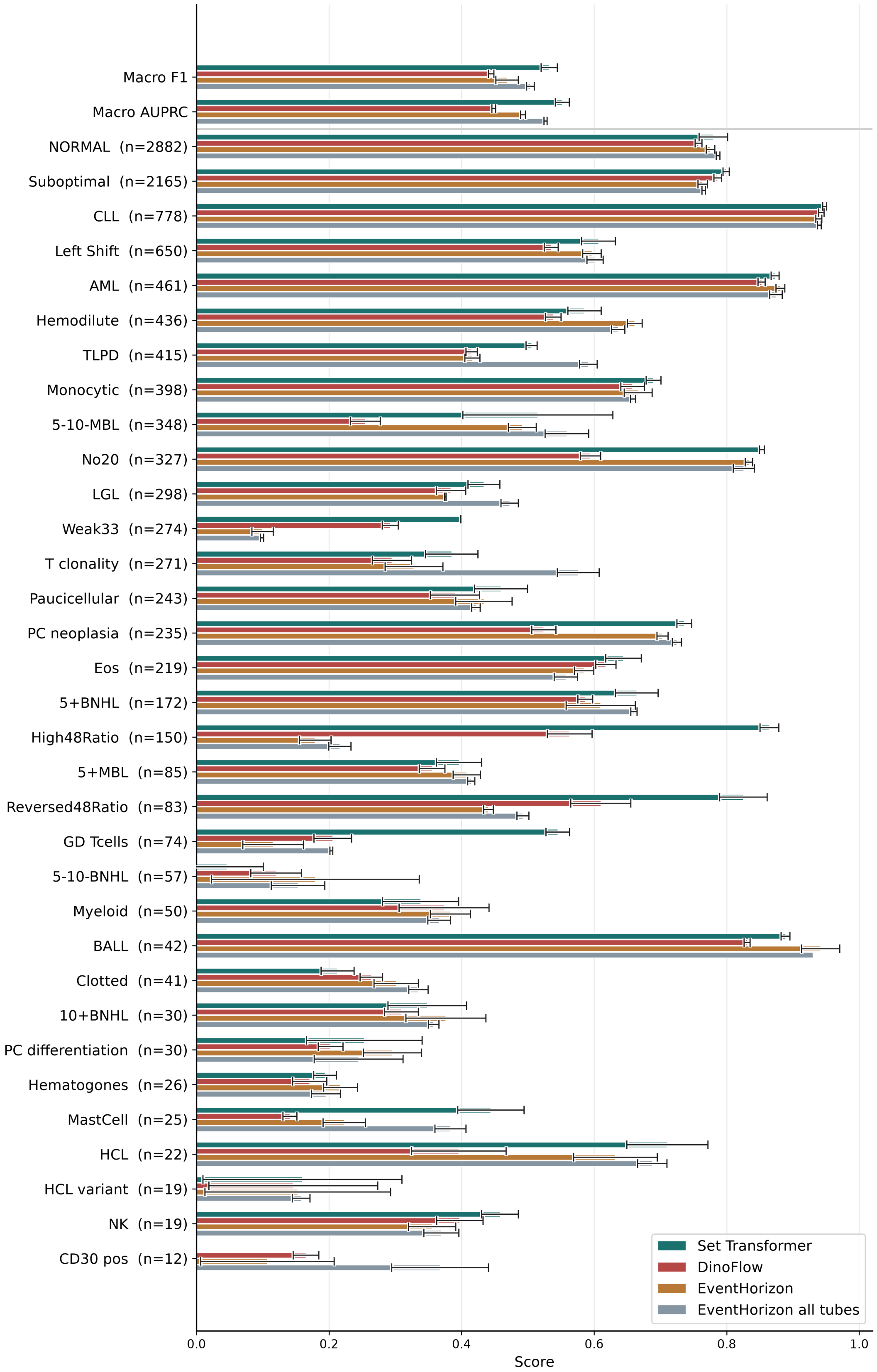

**Fig. S2**: Binary diagnosis performance on the triage screening evaluation. Macro F1 and macro AUPRC (top), then per-label F1 ordered by holdout prevalence (mean ± SD over three seeds). EventHorizon and DinoFlow use frozen self-supervised specimen embeddings with MLP probes; ST-FPS is an end-to-end supervised baseline. EventHorizon is shown for triage B/T/myeloid tubes only and for all available tubes per specimen.

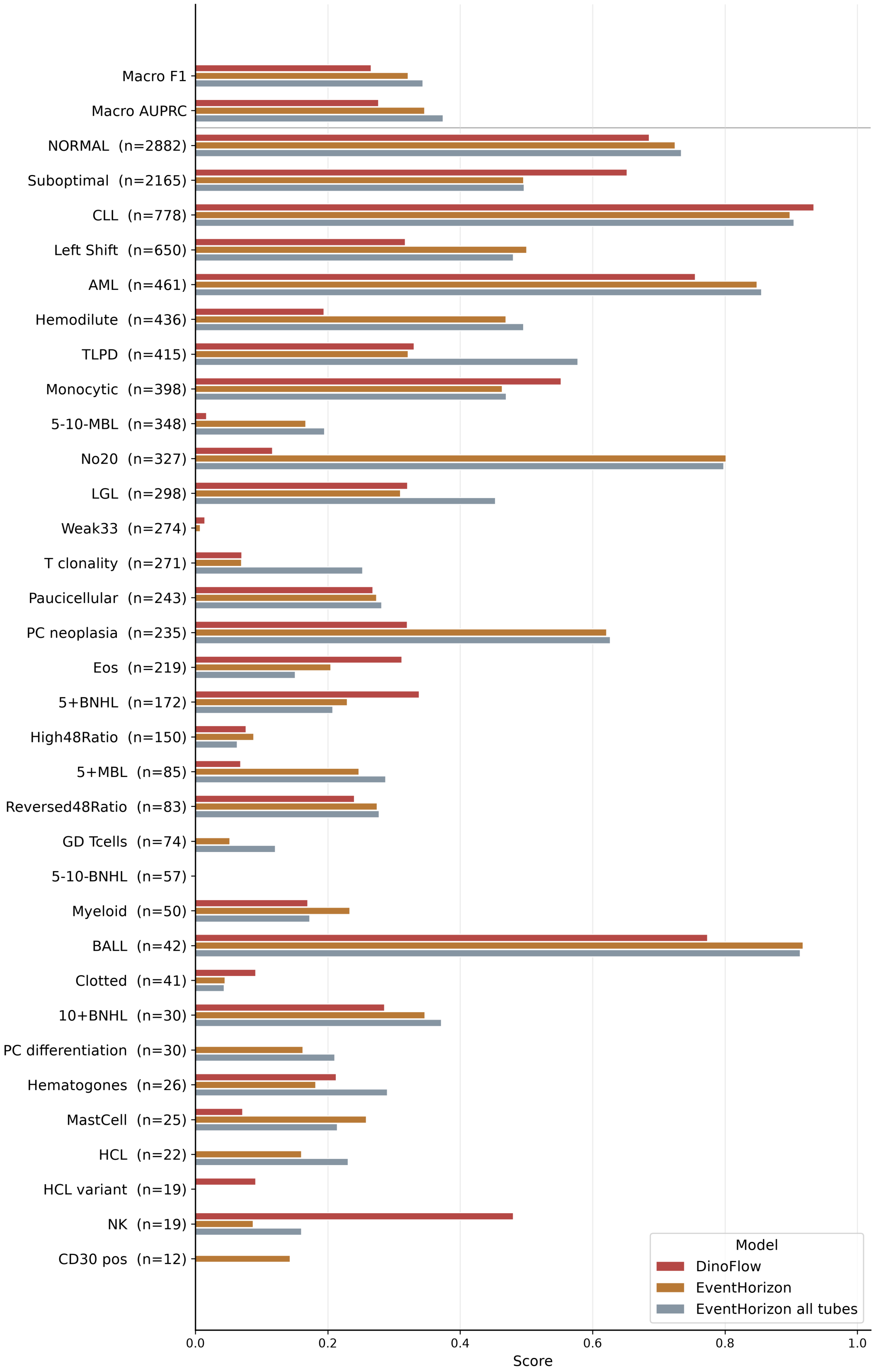

**Fig. S3**: Same layout as Figure S2 with 5-NN probes.

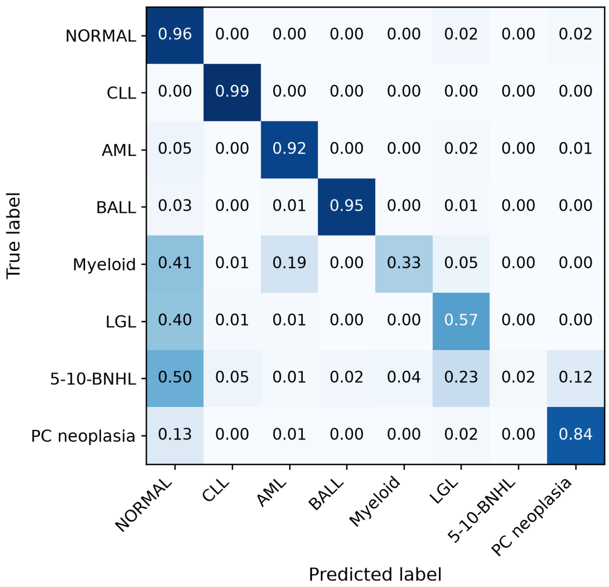

(a) Set Transformer

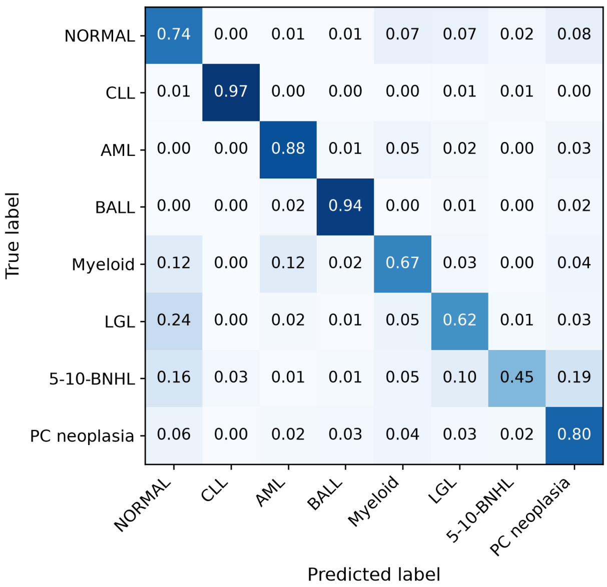

(b) DinoFlow

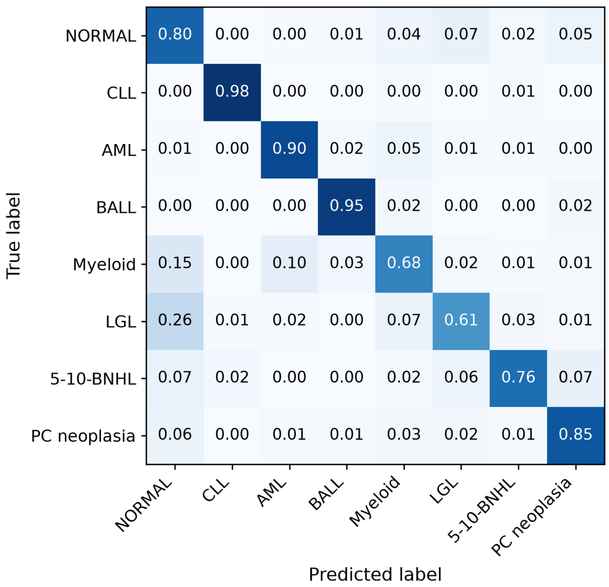

(c) EventHorizon (Triage-Only)

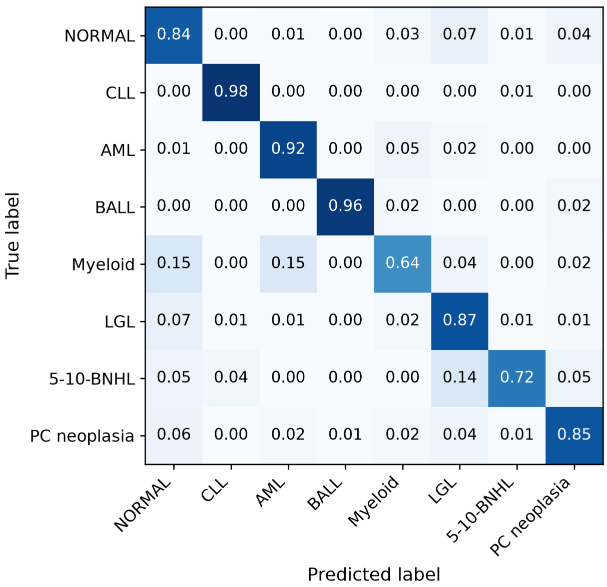

(d) EventHorizon (All-Tubes)

**Fig. S4**: Row-normalized confusion matrices on 2026 holdout specimens with exactly one gold label among eight diagnoses ($n = 4\,729$; NORMAL, CLL, AML, BALL, Myeloid, LGL, 5-10-BNHL, plasma-cell neoplasia). Each panel uses the same binary models as the main comparison (ST-FPS; MLP probes on DinoFlow, EventHorizon triage-only, and EventHorizon all-tubes embeddings). The predicted class is the arg max over the eight probabilities. Counts are pooled over three seeds. Rows are true labels; columns are predicted labels; cell values are the fraction of that true class assigned to each prediction.

#### 2.5 Biological and technical organization of the latent space

The following analyses complement Section 3.10 by evaluating the sensitivity of the graph-based results to graph-construction and community-detection parameters. We repeated the modularity analyses using $k = 5$, 10, and 30 nearest neighbors, and evaluated Leiden community purity across the same neighborhood sizes and resolutions 0.5, 1.0, and 2.0. The main qualitative conclusions were consistent across these parameter settings.

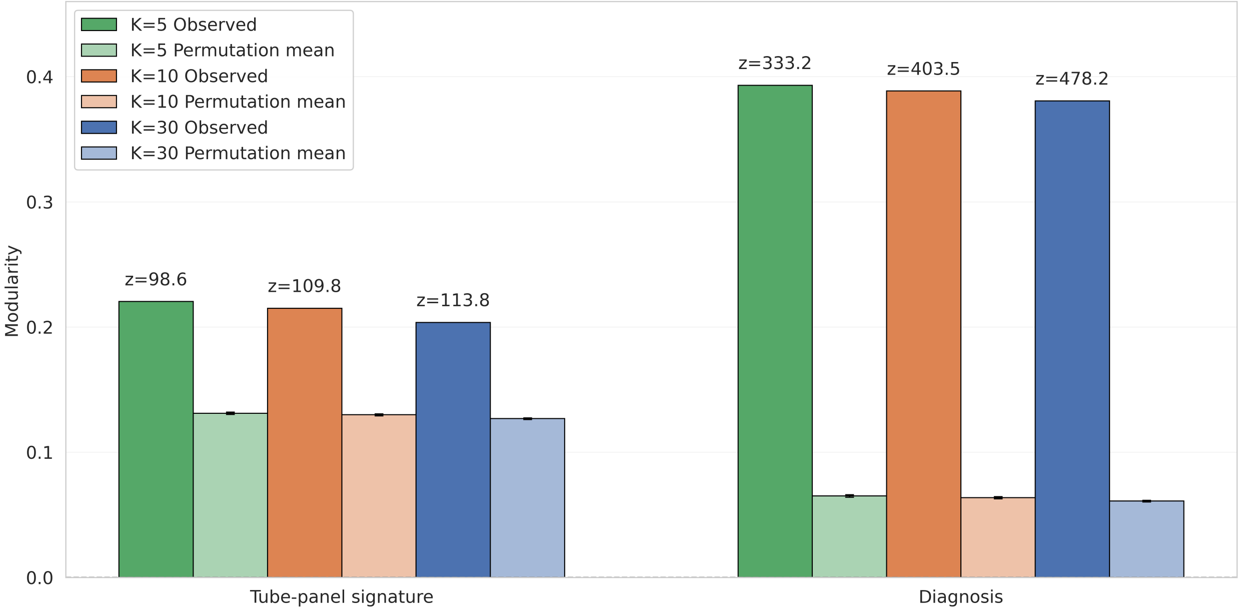

**Fig. S5**: Specimen-level graph modularity analysis of EventHorizon embeddings generated from all tubes available for each specimen. kNN graphs were constructed using $k = 5$, $k = 10$, and $k = 30$. Observed modularity was evaluated with respect to diagnosis and tube-panel signature and compared against stratified permutation means. Text annotations above the observed bars indicate the corresponding permutation-derived $z$-scores, showing the degree of enrichment relative to the null expectation.

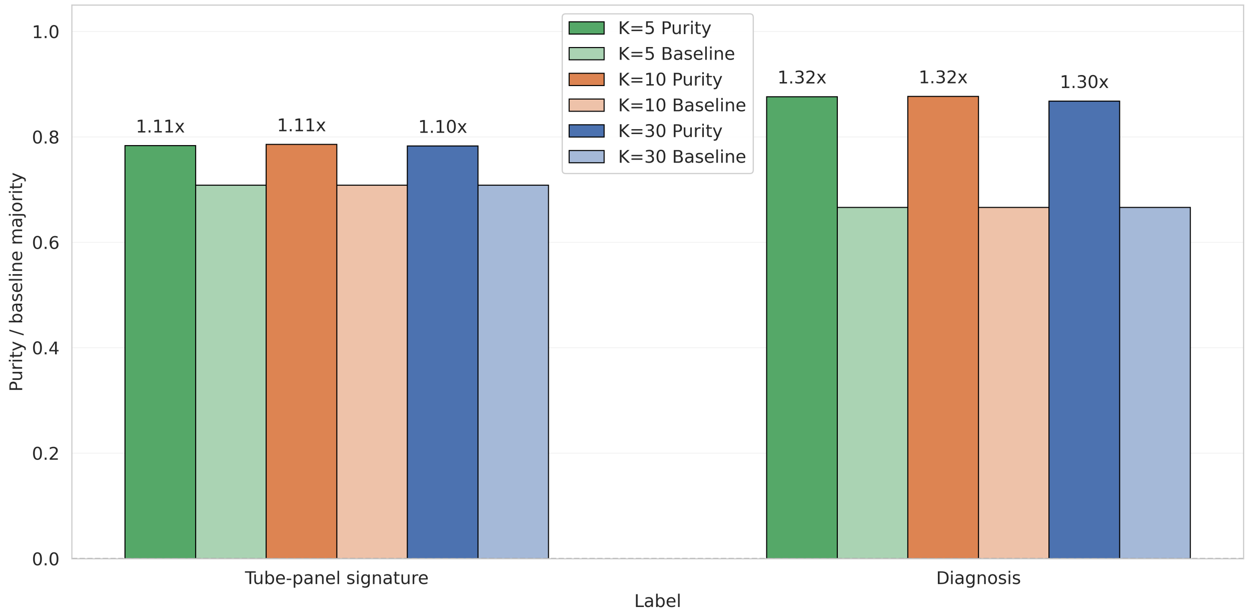

**Fig. S6**: Specimen-level Leiden community purity analysis at resolution 0.5 of EventHorizon embeddings generated from all tubes available for each specimen. kNN graphs were constructed using $k = 5$, $k = 10$, and $k = 30$. Weighted community purity was evaluated with respect to diagnosis and tube-panel signature and compared against majority-label baselines. Text annotations above the purity bars indicate the fold enrichment of community purity relative to the corresponding global majority-label baseline.

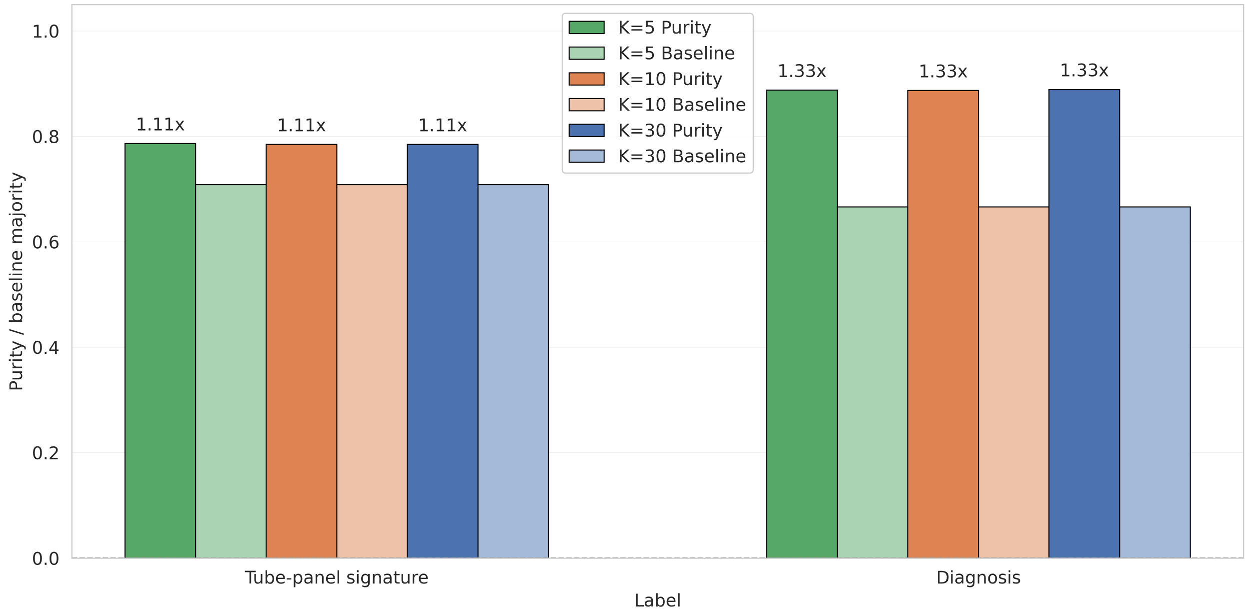

**Fig. S7**: Specimen-level Leiden community purity analysis at resolution 1.0 of EventHorizon embeddings generated from all tubes available for each specimen. kNN graphs were constructed using $k = 5$, $k = 10$, and $k = 30$. Weighted community purity was evaluated with respect to diagnosis and tube-panel signature and compared against majority-label baselines. Text annotations above the purity bars indicate the fold enrichment of community purity relative to the corresponding global majority-label baseline.

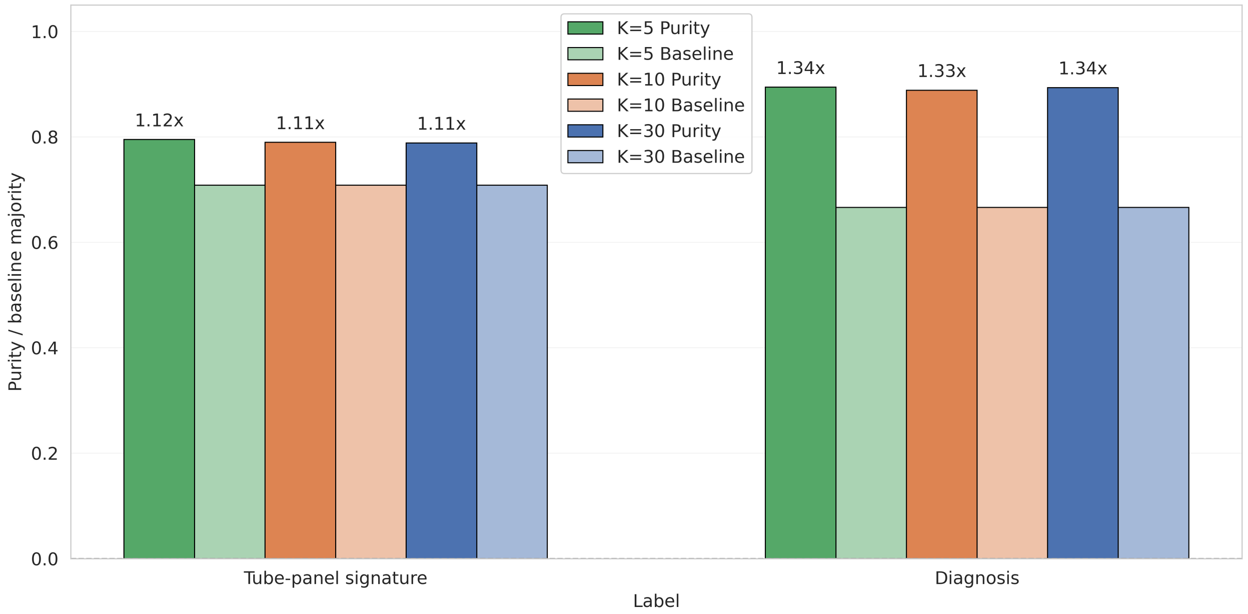

**Fig. S8**: Specimen-level Leiden community purity analysis at resolution 2.0 of EventHorizon embeddings generated from all tubes available for each specimen. kNN graphs were constructed using $k = 5$, $k = 10$, and $k = 30$. Weighted community purity was evaluated with respect to diagnosis and tube-panel signature and compared against majority-label baselines. Text annotations above the purity bars indicate the fold enrichment of community purity relative to the corresponding global majority-label baseline.

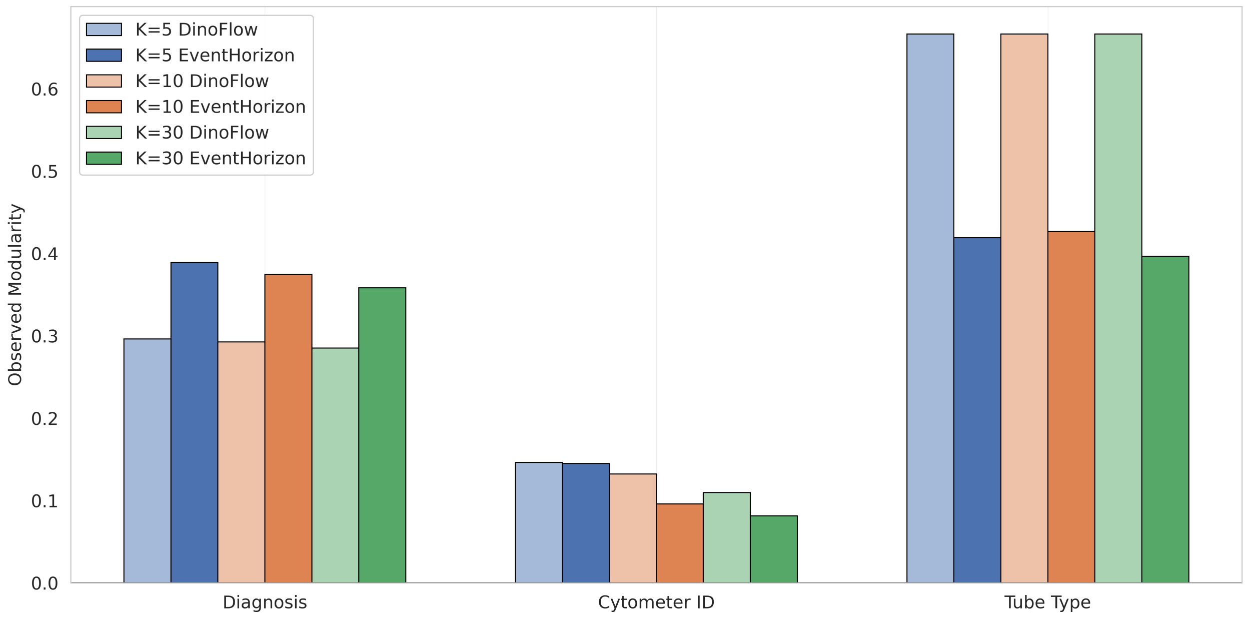

**Fig. S9**: Effect of kNN neighborhood size on label-associated graph modularity. EventHorizon and DinoFlow embeddings were computed for individual B-cell, T-cell, and myeloid triage tubes, with each node representing one tube-level embedding. kNN graphs were constructed with $k = 5$, $k = 10$, and $k = 30$, and observed modularity was computed with respect to diagnosis, cytometer ID, and tube type. Paired bars compare the two embedding models for each label and neighborhood size.

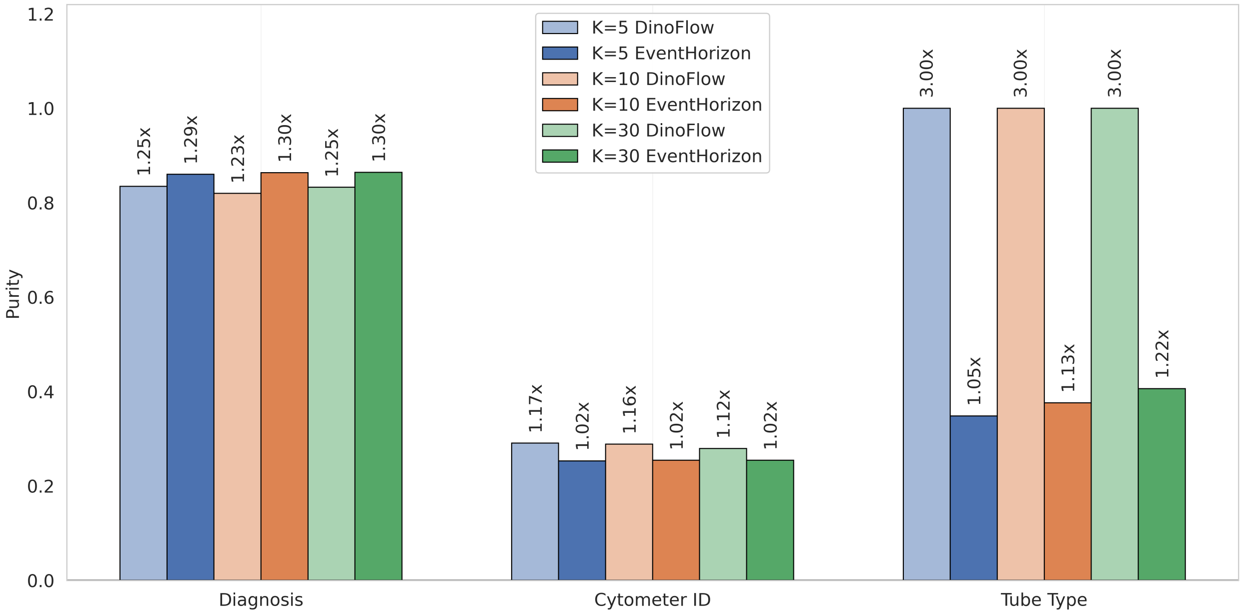

**Fig. S10**: Effect of kNN neighborhood size on Leiden community purity. EventHorizon and DinoFlow embeddings were computed for individual B-cell, T-cell, and myeloid triage tubes, with each node representing one tube-level embedding. kNN graphs were constructed with $k = 5$, $k = 10$, and $k = 30$, followed by Leiden clustering at resolution 1.0. Bars show weighted mean community purity with respect to diagnosis, cytometer ID, and tube type. Text annotations indicate the fold enrichment of purity relative to the corresponding global majority-label baseline.

#### 2.6 Additional evaluation on the immature T add-on panel

| Specimen Type | Counts |
|---|---|
| Peripheral Blood | 536 |
| Bone Marrow | 275 |
| Lymph Node | 182 |
| Other Tissue | 72 |
| Mediastinum | 72 |
| Other Fluid | 50 |
| Thymus | 17 |
| CSF | 4 |
| Total | 1208 |

| Diagnosis | Counts |
|---|---|
| Other Neoplasm | 278 |
| Normal | 246 |
| TCUS† | 210 |
| TLPD† | 197 |
| T-LBL | 181 |
| Thymoma/Thymic Hyperplasia | 96 |
| Total | 1208 |

**Table S7**: Summary of specimens and diagnoses for the immature T add-on evaluation task. †In this context we use TLPD as a shorthand label that subsumes all other T-cell neoplasms besides T-LBL, mostly mature neoplasms, and TCUS as a shorthand for any non-neoplastic expansions of T-cell populations, including T-cell clones of undetermined significance, non-clonal expansions of large granular lymphocytes, etc. "Other neoplasms" include any non-T-lineage neoplasms, which are mostly other acute leukemias. Note that cases where the immature T tube had fewer than 16 384 measured events are excluded from inference, hence the total number of specimens is smaller than shown in Table S1. T-LBL: T-cell lymphoblastic leukemia/lymphoma.

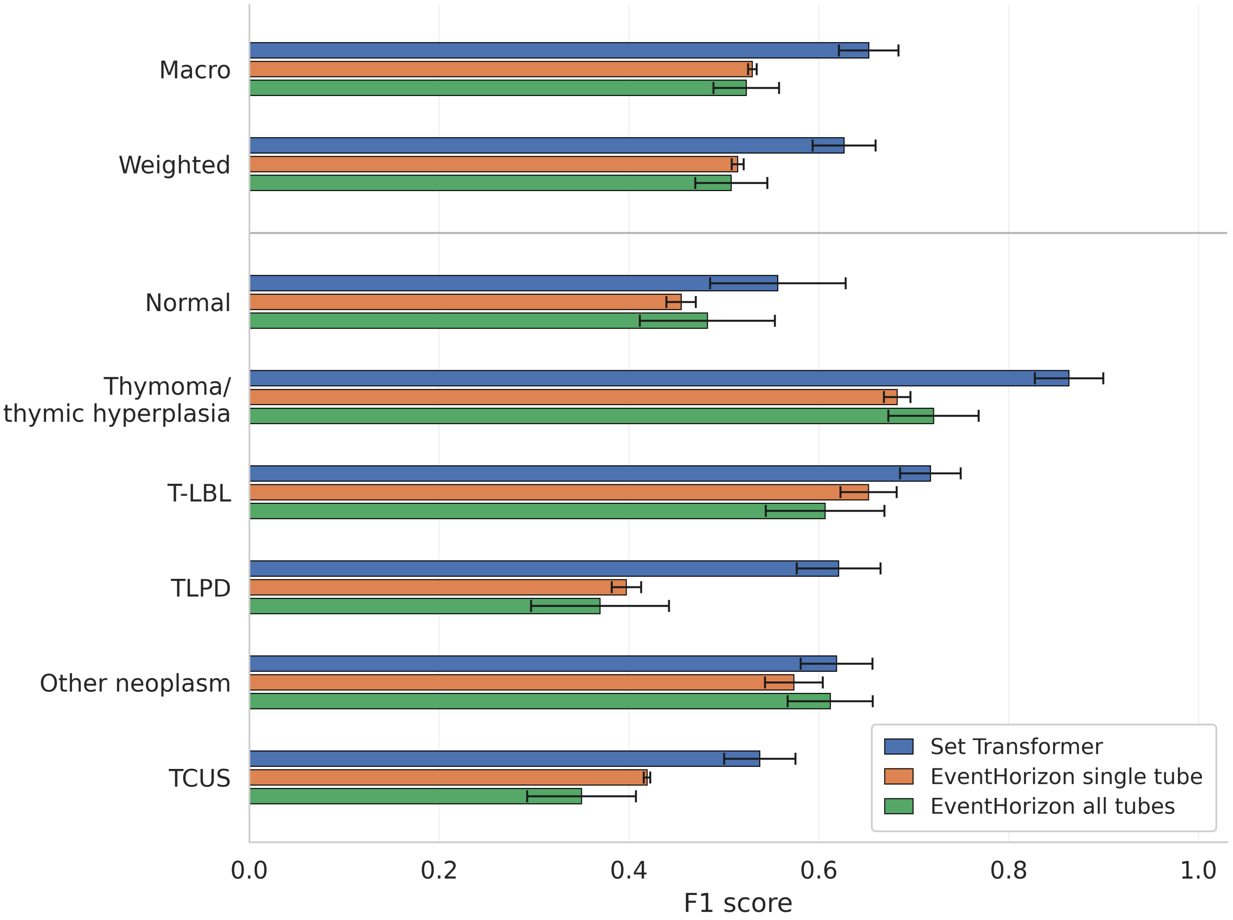

**Fig. S11**: Comparison of EventHorizon embeddings with a supervised ST-FPS baseline on the immature T add-on classification task. Per-class and aggregate F1 scores are shown for k-nearest ($k = 5$) neighbors classifiers trained on frozen EventHorizon embeddings generated from either the immature T add-on tube alone or all available tubes from each specimen. Performance is shown alongside ST-FPS, a supervised transformer-based model trained directly on the same diagnostic classification task.

As an additional evaluation of panel flexibility, we assessed EventHorizon on the immature T add-on panel, a rare, clinically specialized panel with limited representation during pre-training. The specimen and class distributions are summarized in Table S7. We generated embeddings using (1) the immature T add-on tube alone with 16 384 events and (2) all available tubes per specimen with 32 768 events split across tubes. kNN classifiers were trained on both embedding types, and performance was compared to a supervised ST-FPS baseline. Figure S11 shows the resulting per-class F1 scores. The ST-FPS model achieved higher performance across several classes, particularly for TCUS, TLPD, and normal categories. To assess embedding structure, we applied UMAP to the immature T embeddings (Figure S12). The embeddings separate strongly by specimen type. Within specimen type clusters, specimens are further organized by diagnosis. A small, dense cluster corresponding to repeated control specimens is also observed, indicating reproducible embeddings across technical replicates.

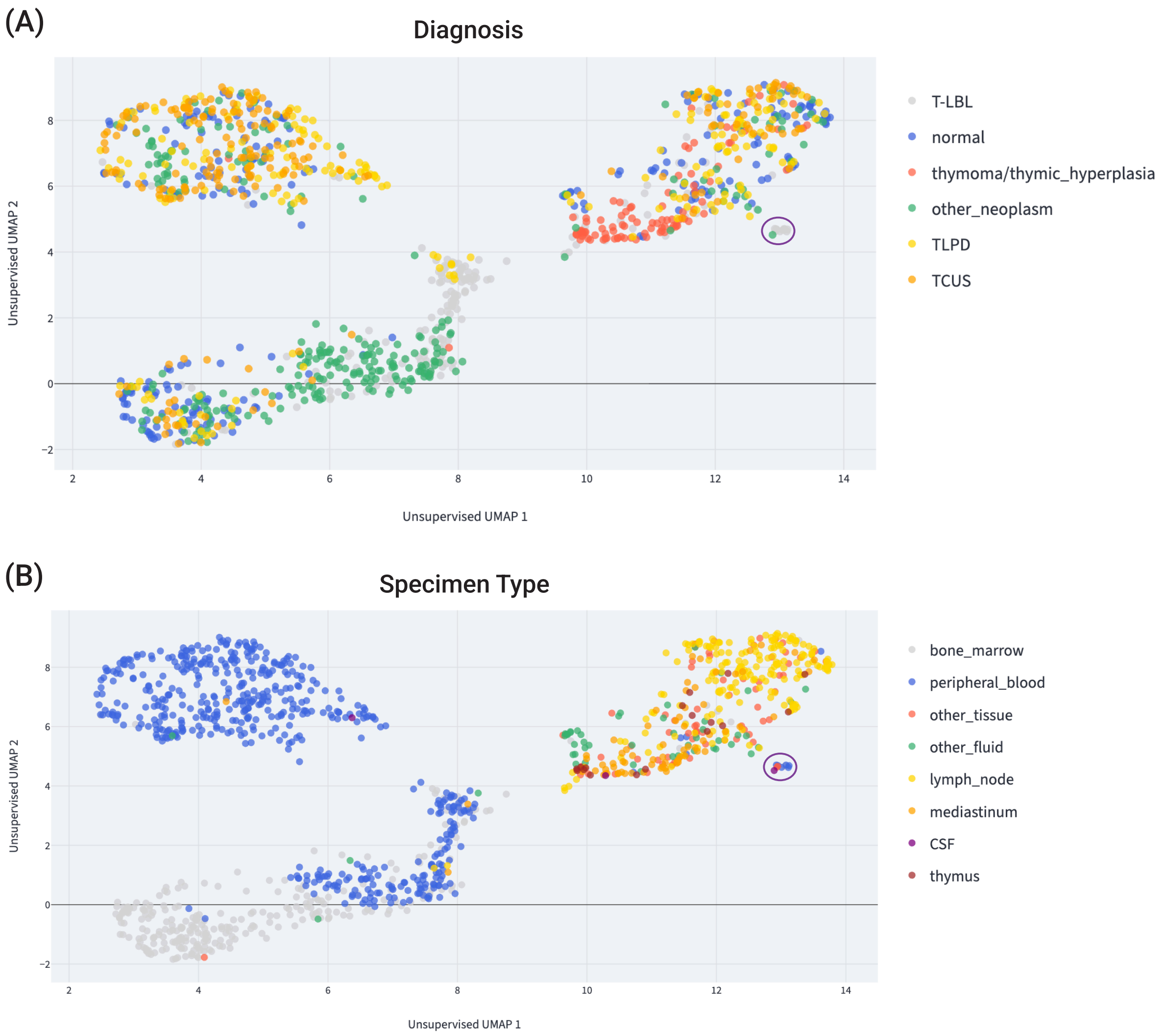

**Fig. S12**: UMAP [27] of EventHorizon embeddings for cases in our dataset for which an immature T add-on was run (see Table S1 and Table S3). Embeddings were generated using only the immature T add-on panel. The circled cluster (in purple) contains CAP rare antigen samples of a CD1a expressing cell line (RFAV1) distributed over the course of 5 years, demonstrating good reproducibility across time, along with a single clinical case containing 97% immature T cells.

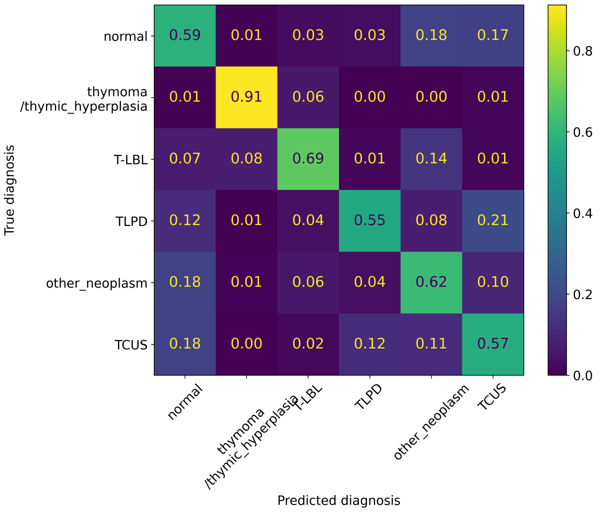

(a) Set Transformer

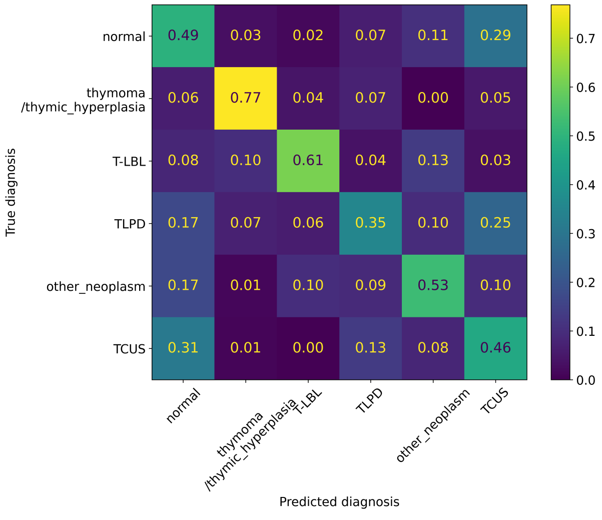

(b) EventHorizon (Single-Tube)

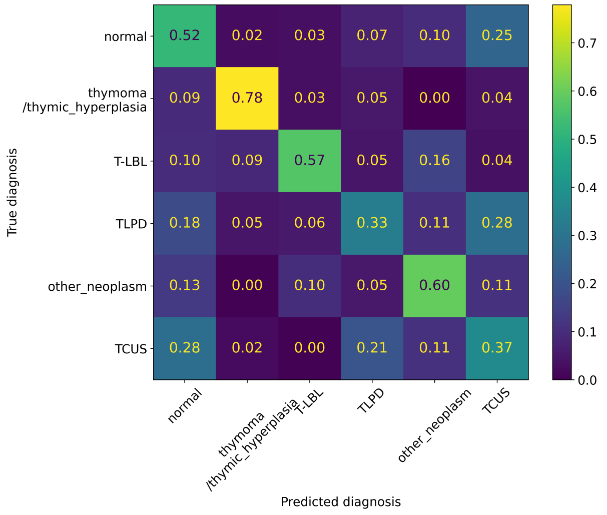

(c) EventHorizon (All-Tubes)

**Fig. S13**: Normalized confusion matrices for the immature T add-on classification task. Panels show (a) ST-FPS single-tube, (b) EventHorizon single-tube, and (c) EventHorizon all-tubes embeddings.

#### 2.7 Additional evaluation on cytoplasmic add-on panels

| Specimen Type | Counts |
|---|---|
| Peripheral Blood | 1782 |
| Bone Marrow | 1343 |
| Lymph Node | 149 |
| Other Tissue | 83 |
| Mediastinum | 54 |
| Other Fluid | 53 |
| Thymus | 11 |
| CSF | 9 |
| Total | 3484 |

| Diagnosis | Counts |
|---|---|
| Myeloid | 2233 |
| B-cell | 649 |
| T-cell | 324 |
| Normal | 202 |
| B/Myeloid | 39 |
| T/Myeloid | 37 |
| Total | 3484 |

**Table S8**: Summary of specimens and diagnoses for the cytoplasmic add-on evaluation task. Note that cases where the cytoplasmic add-on had fewer than 16 384 measured events are excluded from inference, hence the total number of specimens is smaller than the combined total of cytoplasmic 1 and cytoplasmic 2 shown in Table S1.

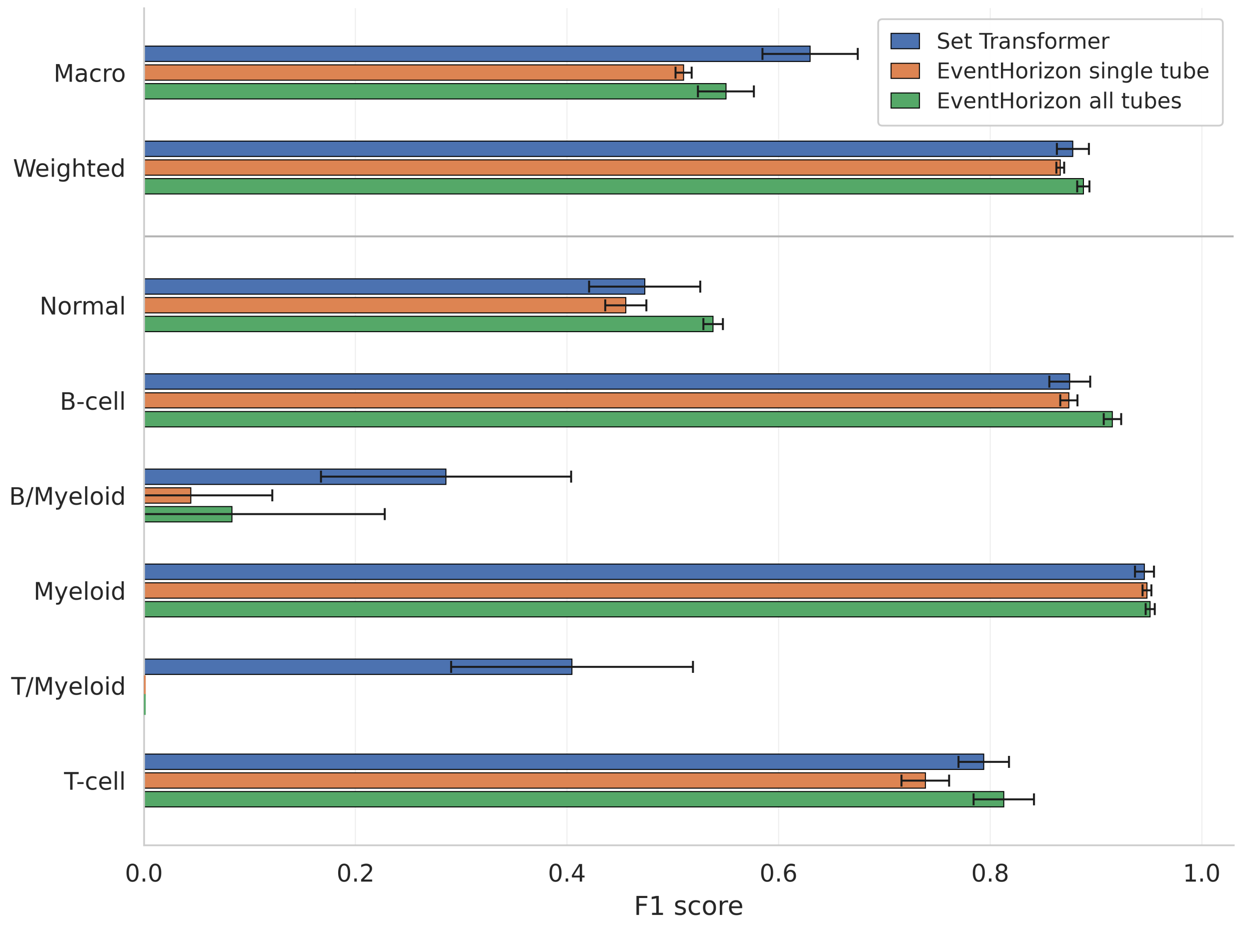

**Fig. S14**: Comparison of EventHorizon embeddings with a supervised ST-FPS baseline on the cytoplasmic add-on classification task. Per-class and aggregate F1 scores are shown for k-nearest neighbors ($k = 5$) classifiers trained on frozen EventHorizon embeddings generated from either the cytoplasmic add-on tube alone or all available tubes from each specimen. ST-FPS is included as a supervised comparison model trained directly on the cytoplasmic add-on diagnostic labels.

As a second additional evaluation of panel flexibility, we assessed EventHorizon on cytoplasmic add-on panels, which have distinct marker compositions and support lineage-focused diagnostic tasks. Rare classes with extremely low counts were excluded, resulting in the distribution shown in Table S8. Embeddings were generated using either (1) cytoplasmic add-on tubes only with 16 384 events or (2) all available tubes per specimen with 32 768 events split across tubes. kNN classifiers were trained on each embedding type and compared to a supervised ST-FPS model. Figure S14 shows per-class F1 scores. Performance varies across classes, with reduced performance observed for mixed-lineage classes, which are underrepresented in the dataset. For the more common single-lineage categories (B-cell, T-cell, myeloid, and normal), kNN classification performance is comparable to the supervised baseline. UMAP visualization of the embeddings (Figure S15) shows separation of specimens by specimen type, with bone marrow and peripheral blood forming distinct clusters from tissue and fluid specimens. Within these clusters, single-lineage diagnoses are broadly separated. Mixed-lineage cases are observed within regions corresponding to related single-lineage categories.

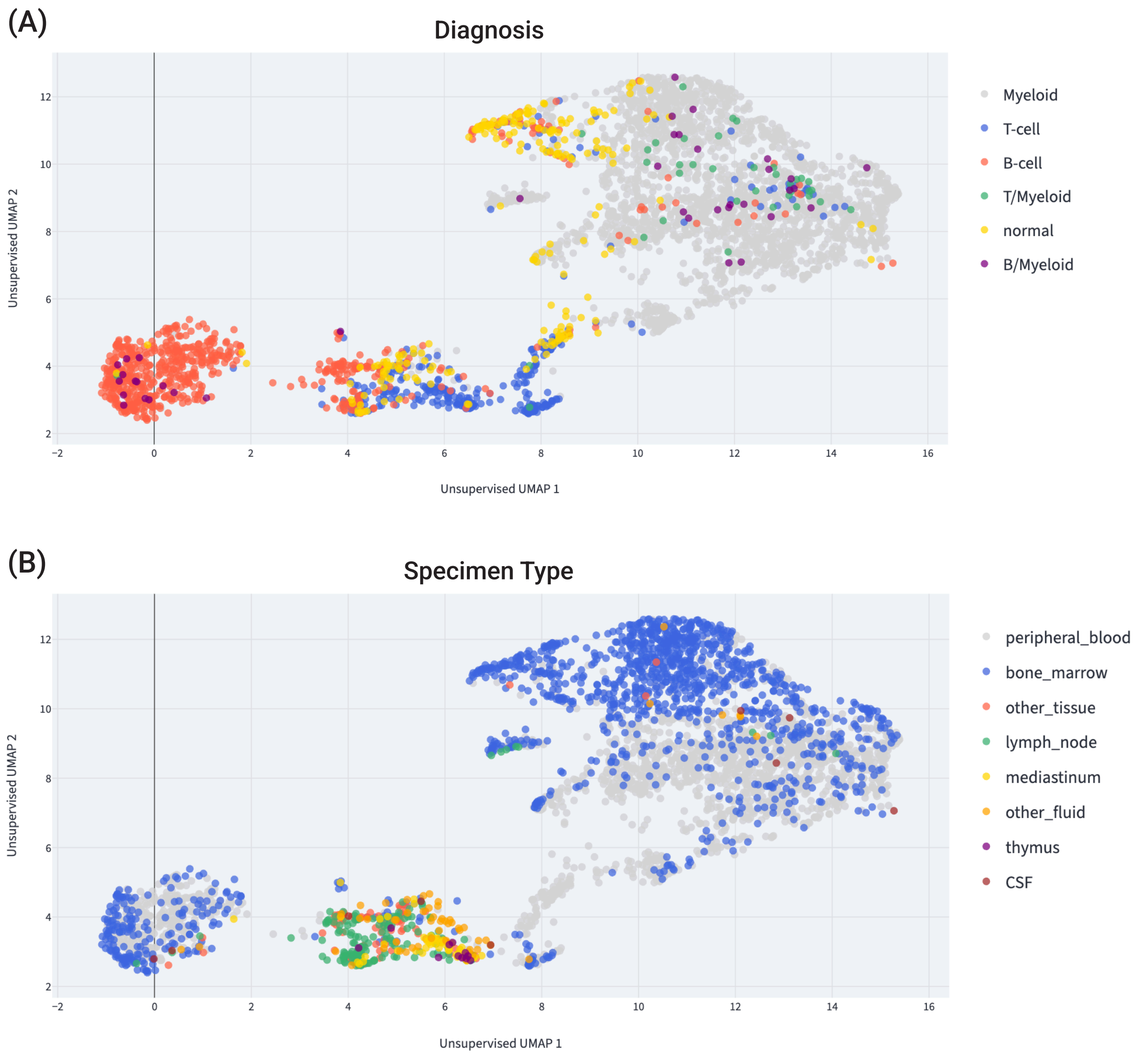

**Fig. S15**: UMAP [27] of EventHorizon embeddings for cases in our dataset for which a cytoplasmic add-on was run (see Table S1 and Table S3). Embeddings were generated using *only* the cytoplasmic add-on panel.

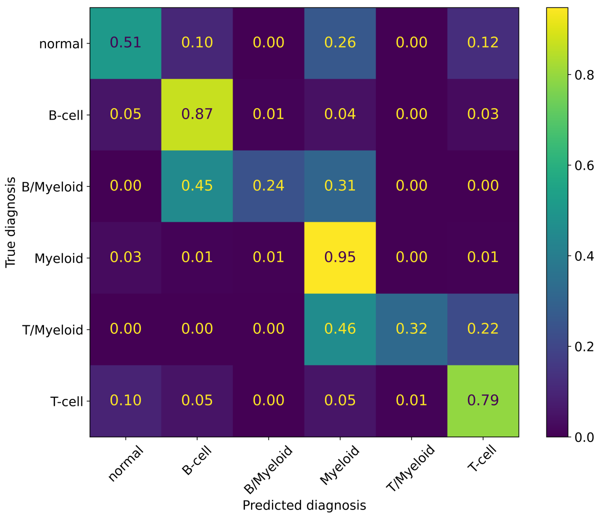

(a) Set Transformer

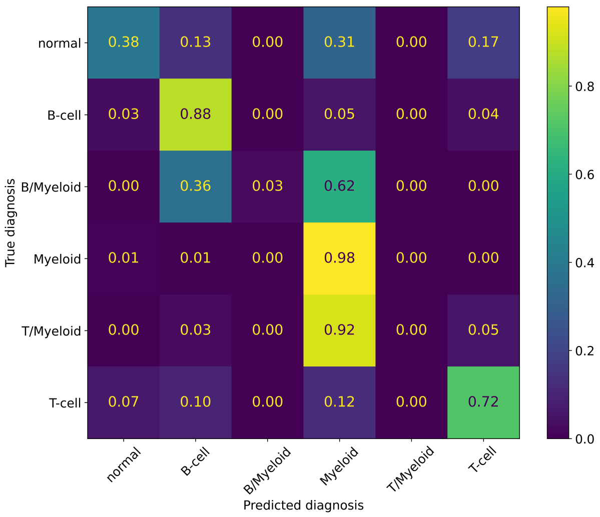

(b) EventHorizon (Single-Tube)

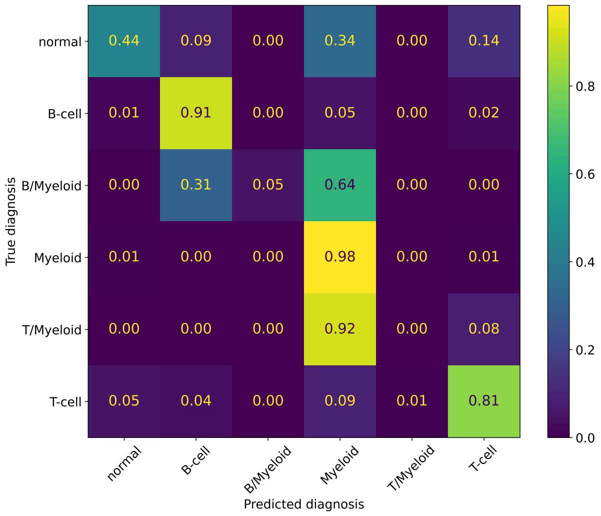

(c) EventHorizon (All-Tubes)

**Fig. S16**: Normalized confusion matrices for the cytoplasmic add-on classification task. Panels show (a) ST-FPS single-tube, (b) EventHorizon single-tube, and (c) EventHorizon all-tubes embeddings.
